# A Systematic Investigation of Overfitting in Maximum Likelihood Phylogenetic Inference

**DOI:** 10.1101/2025.10.07.680876

**Authors:** Anastasis Togkousidis, Olivier Gascuel, Alexandros Stamatakis

## Abstract

Maximum Likelihood (ML) tree inference reconstructs phylogenies from Multiple Sequence Alignments (MSAs). Since MSAs are inherently noisy, ML tools may experience overfitting, whereby the inferred topology incorrectly models noise alongside the true phylogenetic signal. We statistically assess overfitting in ML tools using log-likelihood scores on unseen sites as primary metric. We deploy a 10-fold Monte Carlo cross-validation approach, partitioning 9,062 empirical and 6,342 simulated MSAs into training (80%) and testing (20%) sites. We conduct inferences using RAxML-NG, IQ-TREE, Fast-Tree, and RAxML-NG ES (a recently released Early Stopping version) on the training MSAs. We store all intermediate improved topologies and subsequently evaluate them on the testing sites. We perform a linear regression on the final segments (we use four distinct segment-window configurations) of the derived testing curves, and statistically evaluate the line slopes via the sign test. Our results indicate that ML tools do not overfit. For RAxML-NG (standard and ES) and IQ-TREE, the overall trend is non-significant for 86-98% of the empirical MSAs (across all four segment-window configurations), while for less than 1% the tools exhibit over-fitting. FastTree shows more positive trends, suggesting premature termination, especially on protein MSAs. Topological accuracy curves on simulated data further confirm that tools do not systematically diverge from the true topology. To complement these findings, we test whether strategies to mitigate overfitting can benefit ML inferences. To this end, we also benchmark a site-based holdout validation (HV) version of RAxML-NG. The results confirm that the overfitting is absent and also indicate that excluding MSA sites substantially reduces phylogenetic signal as well as accuracy.

## 1. Introduction

In machine learning, the term *training* refers to the process of estimating the parameters of a statistical model, such that they optimally fit the input data according to a given criterion/loss-function. For instance, in deep neural networks (DNNs), training typically involves iterative methods (Rumelhart et al., 1986), which efficiently adjust the DNN weights (parameters) to progressively minimize prediction errors. *Overfitting* occurs when the model that is being trained does not solely capture the underlying pattern, but also the inherent noise in the data (Geman et al., 1992). It is often a consequence of *over-training* an *over-parameterized* model (see Figure 1), relative to the size and the intrinsic properties of the input data (Bishop and Nasrabadi, 2006; Hinton et al., 2012). However, extensive training of the model, despite strong indications of convergence in the training score/error, does not necessarily induce overfitting (Zhang et al., 2016; Belkin et al., 2019). Whether or not overfitting occurs heavily depends on the statistical model and the specific dataset being analyzed.

**Figure 1.**
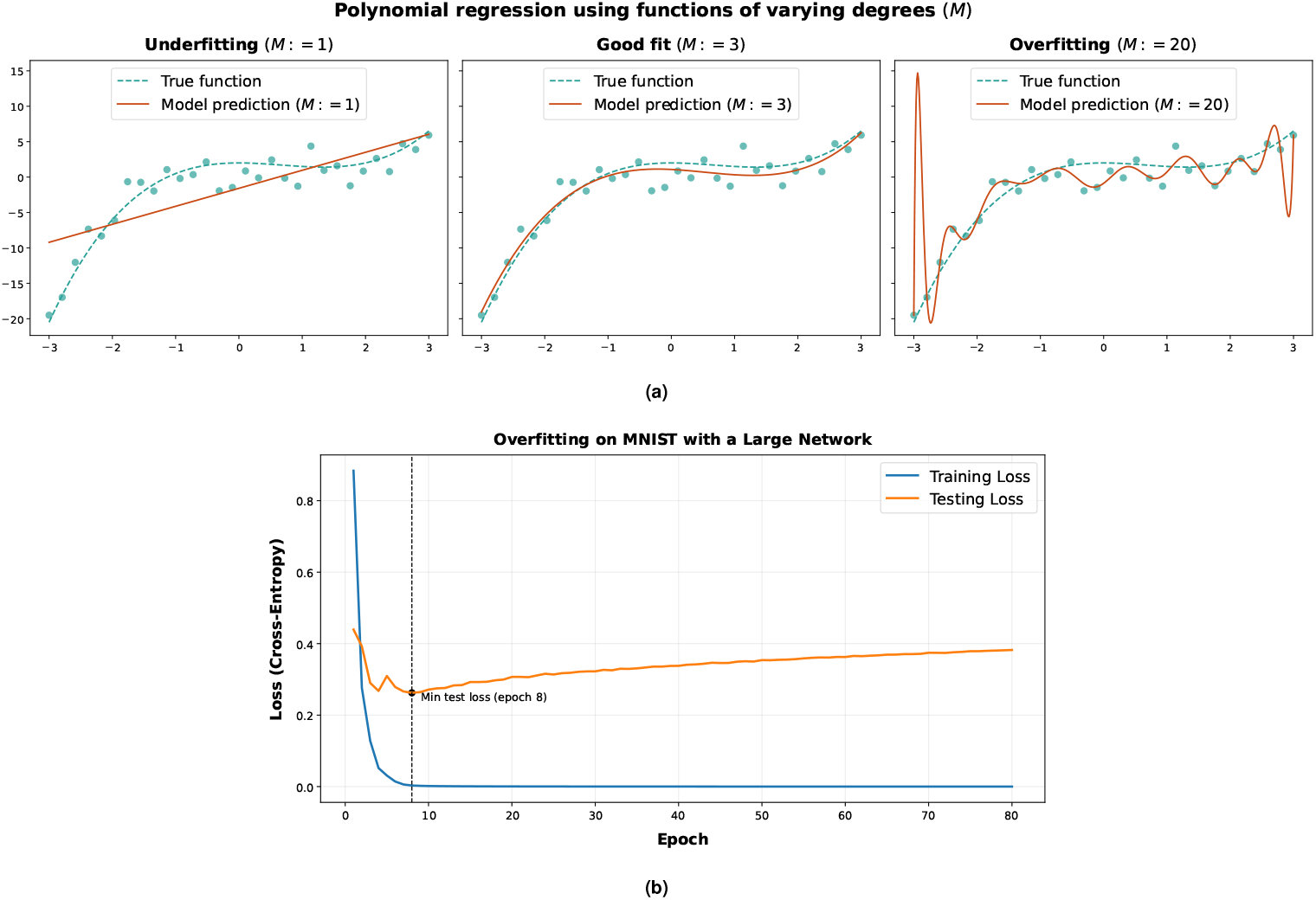
(a) Numerical examples of underfitting, a good fit, and overfitting in polynomial regression. The example shows how intrinsic noise in the data may induce overfitting under parameter-rich models. The data points were generated via a cubic function (indicated as “True function” in the Figure legends) to which we injected Gaussian noise. We regress the input data using three polynomial functions of varying degrees (*M*). The linear model (*M* := 1) fails to capture the nonlinear data pattern and therefore underfits. The cubic model (*M* := 3) effectively captures the true pattern and yields a good fit. The parameter-rich model (*M* := 20) incorrectly captures noise alongside the true signal, resulting in overfitting. **(b) Numerical example of overfitting emerging during the training of a parameter-rich DNN**. The plot corresponds to empirically derived learning curves obtained from training a DNN on a small random subsample (1,600 and 400 data points for training and testing, respectively) of the MNIST dataset (LeCun et al., 2002). The DNN has four fully connected layers with widths of 784, 1024, 1024, and 10. We train the DNN via the Adam optimizer (Kingma and Ba, 2017) but disable any form of regularization or early stopping. After iteration/epoch 8, the testing loss function progressively increases and diverges from the training one, which continues to decrease and eventually attains zero loss.

Phylogenetic tree inference aims to identify the unrooted binary tree that best describes the evolutionary relationships among the sequence data in a given Multiple Sequence Alignment (MSA). Identifying this tree translates into a computational optimization problem under the Parsimony (Fitch, 1971) and Likelihood (Felsenstein, 1981) criteria. Due to the prohibitive computational complexity of Maximum Parsimony (MP) and Maximum Likelihood (ML) based tree inference (Day et al., 1986; Roch, 2006), contemporary software tools deploy ad hoc heuristics to identify trees with “good” optimality scores that typically represent local optima. Under ML, the model comprises the binary tree topology, its branch lengths, and the substitution model parameters, which jointly describe the stochastic process under which the observed sites in the MSA evolve. Heuristics strive to estimate the statistical model parameters, by optimizing the tree topology (a discrete parameter) and the continuous parameters (branch lengths and the substitution model parameters (Yang, 2014)).

Genomic sequences, however, are subject to noise, that stems from stochastic and/or systematic sources. Stochastic noise reflects the randomness of evolution, resulting in sites that deviate from the expected theoretical distributions (Hillis and Huelsenbeck, 1992). Furthermore, sampling bias (Yang et al., 1995) is superimposed onto stochastic noise, which questions the statistical adequacy of the input MSAs as proxies for the underlying site distribution (Togkousidis et al., 2025). Systematic noise, on the other hand, is more prevalent in empirical MSAs, and is induced by a plethora of phenomena, such as sequencing errors (Moutsopoulos et al., 2021), alignment errors (Dress et al., 2008), and discordant gene histories in phylogenomic MSAs (Maddison, 1997).

A key analogy between ML tree inference heuristics and deep learning methods is that they both employ iterative algorithms to optimize the model-fit to the input data, either by minimizing the error in DNNs, or by maximizing the log-likelihood score under ML. A second analogy is that both DNNs and likelihood-based phylogenetic models (discussed in the next paragraph) constitute parameter-rich models. As shown in the example of Figure 1, parameter-rich models are susceptible to overfitting. In the polynomial regression example of Figure 1a, overfitting occurs with a high degree polynomial (*M* := 20, where *M* indicates the polynomial degree). In contrast, a cubic-model (*M* := 3) suffices to capture the underlying pattern. Under overfitting (*M* := 20), the predictive accuracy on unseen (testing) data is suboptimal, as the model also fits the training data’s noise on top of the true signal. This example directly applies to DNNs, which are used both, for regression, and classification problems. DNNs generally constitute parameter-rich models that comprise thousands or millions of parameters/weights (Montavon et al., 2012). The DNN parameter/weights initially have random values, and they are adjusted via successive training iterations. As shown in Figure 1b, overfitting typically emerges *during* training: after a certain point (iteration/epoch 8 in the example), the DNN begins to model the intrinsic noise in the training data, and therefore, its performance on unseen (testing) data deteriorates. A common technique to mitigate overfitting in deep learning is to apply holdout-based early stopping (Srivastava et al., 2014). Under this approach, the input dataset is randomly split into model-training and validation data. During training, the model—with the currently inferred weights—is periodically evaluated on the validation set, and optimization halts once the validation error starts to increase.

In analogy, likelihood-based phylogenetic tree models are also parameter-rich. Substitution models typically contain (only) a small number of free parameters (e.g., 9 for GTR+Γ4 (Tavaré, 1986)), and numerous methods/software have been proposed for selecting the best model (Sullivan and Joyce, 2005; Lefort et al., 2017; Darriba et al., 2020) based on the AIC, (Akaike, 1974), or the BIC (Schwarz, 1978) scores. Related work also addresses the risk of over-parameterization in profile mixture models (Baños et al., 2024). The role of the *tree topology*, though, as a potential source of overfitting has received little attention, despite being the most biologically meaningful parameter. The phylogenetic tree involves one parameter/length per branch (e.g., ~2,000 edges in a tree comprising 1,000 tips), plus the tree topology itself that forms part of a vast, super-exponentially large space. ML tree inference, similar to DNN training, proceeds in discrete iterations, which correspond to single topological updates (see below). ML tools initiate tree searches either from a random, or parsimony-based tree, and gradually improve with respect to the branch-lengths and tree topology. The process terminates when they reach a local likelihood optimum. The likelihood is computed on the input MSA, which is analogous to the training set in a DNN.

As a result, phylogenetic models may also become over-parameterized regarding the size, the amount of noise, and intrinsic properties of the input data (Häuser et al., 2025). On the other hand, ML inference and deep learning exhibit a substantial difference. In a fixed DNN graph, model parameters are continuous and typically homogeneous in nature and significance, whereas under ML, the discrete tree topology is biologically more important than the remaining parameters. To our knowledge, relatively few studies have explicitly addressed topological overfitting in ML tree inference (Berry et al., 2000; de Vienne et al., 2017), that is, the tree topology itself as a potential source of overfitting. For instance, Berry et al. (2000) evaluate overfitting within a static framework, by applying post-hoc overfitting-detection criteria to trees that had already been inferred. They argue that weakly supported branches may indicate over-parametrization, and they propose methods to collapse such branches into multifurcations. While this approach to mitigating over-parametrization is useful, it is orthogonal to the focus of our study (see below). We nevertheless revisit the issue of overfitted branches in the Discussion section.

Our work primarily builds on the study by de Vienne et al. (2017), which—although published only as a preprint—served as a key reference point. We believe, however, that this study exhibits certain limitations, which we discuss later in the main text and in the Supplementary Material. Analogous to the approach in de Vienne *et al*., we assume that ML tools operate on fully resolved bifurcating trees, which may indeed be over-parameterized given the input data, and examine the dynamics by which overfitting (potentially) emerges *during* the tree inference process. We assess whether iterative heuristics implemented in ML tools reach a turning point during the optimization process that is determined by the phylogenetic signal-to-noise ratio of the input MSA, beyond which, further iterations merely and increasingly fit noise (as in Figure 1b). We define iterations based on rudimentary updates to the current best-scoring tree topology; that is, each newly accepted bifurcating tree corresponds to a distinct iteration. Our approach resembles dynamic overfitting detection in DNN training, as well as in other parameter-rich models (Figure 1b). On the other hand, because ML tools maximize the log-likelihood score, such a degradation manifests itself as a *decrease* in likelihood on unseen (testing) sites, in contrast to DNNs, where overfitting is observed as an *increase* in prediction error (see Figure 1b). Another major difference is that we optimize in a space that is both discrete (topology) and continuous (branch lengths) (Billera et al., 2001). As a consequence our training/testing curves are inherently “bumpy” (Figure 2) since topological changes may induce abrupt steep likelihood increases. Hence, the curves are less smooth and thus more difficult to analyze.

**Figure 2.**
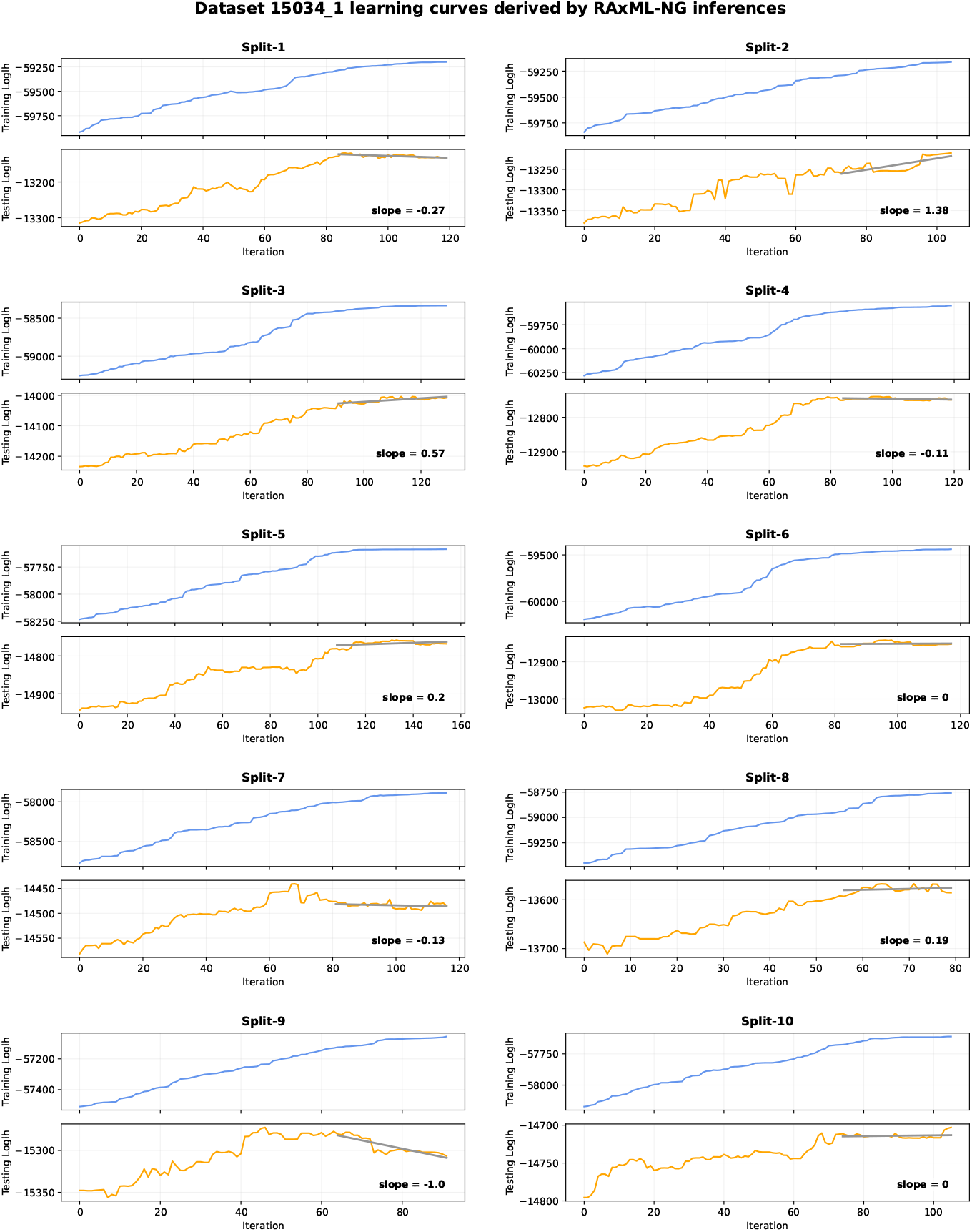
Exemplary learning curves corresponding to the anecdotal empirical DNA dataset 15034_1, comprising 222 taxa, 2,658 sites, and 1,237 patterns. The training log-likelihood curves (upper panel of each split-plot) increase monotonically and are comparatively smooth. The testing log-likelihood curves (lower panel) may be more “bumpy” and not necessarily monotonic. To capture the testing curves’ behavior towards the end of the inference, we perform linear regression on the final curve-segments (here, *n* = 30%*N*). The fitted regression lines are shown in gray color, and the corresponding slope values are reported in the bottom-right corner of each testing log-likelihood panel. Based on the slope values, the final part of the testing curves may exhibit an increasing tendency (splits 2, 3, 5 and 8), form noisy plateaux (splits 6 and 10), or diverge from the curve’s global maximum (splits 1, 4, 7, and 9). Overall, the sign-test yields a non-significant slope trend. In some cases, the fitted line may not fully capture the real testing curve dynamics. For instance, in split 7 the testing curve optimum appears too early, and the regression line gives a near-zero inflection (although, as reported, for *n* = 30%*N* the inferred slope is negative and successfully reflects the model-fit decrease; yet, for the same dataset-split and for *n* = 20%, the estimated slope is 0). Analogously, in split 10 a late increase over the final few iterations is not captured by the fitted line, which yields a zero slope. In all 10 splits, the final, and the optimal tree of the testing curve are statistically indistinguishable, when evaluated via the approximately unbiased (AU) test on the corresponding testing sites (see Discussion).

We address overfitting via two complementary approaches. In the first part, we investigate systematic overfitting trends in three widely used phylogenetic inference tools: RAxML-NG (Kozlov et al., 2019), IQ-TREE (Minh et al., 2020), and FastTree (Price et al., 2010). In this comparison we also include our recently released Early Stopping (ES) version of RAxML-NG (Togkousidis et al., 2025). The term “systematic” indicates that we specifically investigate whether overfitting constitutes *the* statistically expected behavior of ML tools. This question is most relevant for RAxML-NG and IQ-TREE, which deploy thorough tree search heuristics, whereas FastTree and RAxML-NG ES constitute faster alternatives. The main outcome of this analysis is that topological overfitting does not constitute a major issue in any of these tools.

The second part aims to confirm the conclusions of the first, and reinforces our findings. In this part, we benchmark a site-based holdout validation (HV) version of RAxML-NG that is analogous to approaches employed in deep learning. HV shows inferior performance relative to standard RAxML-NG, which conducts inferences on the entire MSA, even on long MSAs. This result is consistent with the findings from the first part of our study, namely that topological overfitting does not constitute a major issue. It also shows that the standard overfitting mitigation technique used in deep learning (HV) does not benefit ML-based tree inference.

## 2. Results

### 2.1. Datasets and Substitution Models

To examine overfitting in ML phylogenetic tree inference, we conducted experiments on 9,062 empirical MSAs (8,229 DNA and 833 AA), sampled from TreeBASE (Piel et al., 2009). We applied no specific sampling criterion, beyond retaining all MSAs which successfully completed all preprocessing and pipeline steps (see below). We further included 6,342 simulated DNA MSAs from a prior benchmark analysis (Trost et al., 2024), comprising 5,020 alignments generated under the GTR+Γ substitution model (Tavaré, 1986) and 1,322 under the JC model (Jukes et al., 1969). Since the results between the GTR+Γ- and JC-generated MSAs are highly similar, we only present results for the former (the results under JC are provided in the Supplementary Material). We discuss this similarity later in the text.

We applied a basic preprocessing step to all MSAs. First, we removed duplicated sequences to retain only unique sequences. Second, we removed gap-only MSA sites from empirical datasets. This preprocessing step was necessary to execute all distinct steps in the experimental pipeline (see Supplementary Figure S3). From the cleaned MSAs, we discarded four-taxon alignments, as they are trivial to analyze. The resulting MSAs comprise up to 10^6^ sites (empirical and simulated datasets), and up to 10^3^ and 500 taxa in empirical and simulated MSAs, respectively. The cleaned MSAs cover the entire difficulty score spectrum as predicted by Pythia v1.2 (Haag et al., 2022). The difficulty score quantifies how easy- or hard-to-analyze an MSA is. Further details regarding datasets, MSA dimensions, and difficulty score distributions are provided in the Supplementary Material (Figures S1–S2).

We treated all datasets as unpartitioned MSAs. This decision is straightforward and consistent with the data. All simulated MSAs were generated as single-partition datasets, and the vast majority of TreeBASE MSAs are, indeed, unpartitioned: ~93% of DNA and ~99% of AA MSAs are provided without any partitioning scheme. To conduct ML tree inferences, we used the GTR+Γ model for all empirical/simulated DNA MSAs (including those simulated under the JC model), and the LG+Γ model (Le and Gascuel, 2008) for protein MSAs (empirical only). In the context of this study, restraining the analyses to unpartitioned MSAs, and using standard substitution models provides a controlled setting to assess overfitting.

To benchmark RAxML-NG HV, we sampled 535 long empirical MSAs (430 DNA and 105 AA) from the preprocessed dataset, along with 241 long simulated DNA MSAs under GTR+Γ. Here, “long” refers to MSAs comprising: (a) more than 10,000 sites and a sites-over-taxa ratio exceeding 10 for empirical MSAs; and (b) more than 5,000 sites and a sites-over-taxa ratio exceeding 5 for simulated MSAs. We relaxed the thresholds for simulated MSAs, since relatively few datasets met the original criteria applied to empirical MSA sampling. We chose those thresholds to ensure that the HV version is evaluated under a statistically meaningful setting. That is, in datasets where the number of observations (sites) is large, both in absolute terms and relative to the number of taxa, we ensure that there is a sufficiently strong phylogenetic signal. As expected, the Pythia score distributions for the sampled, long MSAs shifts toward lower values, indicating easier MSAs (see Supplementary Figure S2). All datasets used in this study are publicly available at https://cme.h-its.org/exelixis/material/overfitting_data.tar.gz.

### 2.2. Overfitting Analysis

We provide a schematic overview of our pipeline (described below) in Supplementary Figure S3. We employ a 10-fold Monte-Carlo cross-validation approach (Picard and Cook, 1984) by randomly splitting each em-pirical/simulated MSA into training and testing sites with a splitting ratio of 80%-20%. As is common in statistical modeling, the training set fraction is high (80%), so as to yield results that are representative of the entire MSA. On the contrary, the proportion of testing sites (20%) suffices to reliably estimate the cross-validated, unbiased value of the optimization criterion (i.e., the likelihood of unseen testing sites; see below). Importantly, this experiment is repeated 10 times (10-fold crossvalidation) to statistically assess the significance of the results.

We conduct ML tree inferences on the training sites of each dataset-split using four tools: RAxML-NG, IQ-TREE, FastTree, and RAxML-NG ES. RAxML-NG ES is our recently developed Early Stopping version of RAxML-NG (Togkousidis et al., 2025), which employs the Kishino-Hasegawa (KH) test (Kishino and Hasegawa, 1989) combined with correction for multiple testing (see Methods) to statistically assess significant log-likelihood differences between the two tree topologies *prior to* and *after* each Subtree Prune and Regraft (SPR) round in RAxML-NG. RAxML-NG ES terminates early when improvements are statistically insignificant (see Methods). The ML tree inferences by the four tools mentioned above begin searching tree space from a parsimony starting tree (i.e., a tree with a “good” parsimony score), which we generate with RAxML-NG on the training MSA of the corresponding dataset-split. We use parsimony starting trees because they have relatively good (i.e., non-random), yet still suboptimal, log-likelihood scores (Yang et al., 1995; Guindon et al., 2010a; Stamatakis, 2014). This choice is appropriate as our focus is on the final stages of the optimization. In other words, we intend to assess whether overfitting occurs during the final optimization steps prior to convergence.

In the course of the ML tree inferences, we store *all* intermediate improved topologies that are accepted by each tool. That is, we store topologies that are visited and retained because they improve the overall log-likelihood score. We adopted this comprehensive tree sampling strategy to capture even the smallest topology-induced log-likelihood improvements with respect to the currently best ML tree. This approach does not only increase the resolution of the learning curves. It also allows for assessing and comparing overfitting tendencies among distinct tools and does not rely on arbitrarily recorded tree topology checkpoints. The challenges involved in implementing this fine-grained tree-sampling approach are described in Methods. Due to this setup, the training curves of all tools increase monotonically, and each point corresponds to an improved (w.r.t. its log-likelihood score) intermediate topology (see Figure 2).

Following the ML tree inferences with the four distinct tools, we evaluate the ML score of all sampled topologies via RAxML-NG using the testing MSA to obtain testing curves for each inference and tool. For this evaluation, we optimize branch lengths and evolutionary model parameters from scratch. Thereby, we assess overfitting specifically *with respect to* the tree topology. We focus on the tree topology as parameter of primary biological interest. Our setup, does not assess the convergence dynamics of the sequence of intermediate trees toward some reference best-known tree topology. While we also perform such an analysis in a distinct step, the objective here is different: we investigate the generalizability of each newly introduced topology, by comparing its likelihood on unseen (testing) sites, relative to its predecessor(s).

After constructing the testing curves, we perform a linear regression on their final segments and extract the corresponding slope. We explore four alternative final segment definitions: the last two points, and the final 10%, 20%, or 30% of the points on each curve. In the following, we refer to these segment configurations as *n* = 2, *n* = 10%*N, n* = 20%*N*, and *n* = 30%*N*, where *n* denotes the number of regression points and *N* the total number of points on the curve. Slopes with an absolute value below 0.1 are set to 0, since such small differences can be attributed to numerical errors (Shen et al., 2020; Stelz et al., 2025). We thereby derive 10 slopes per dataset, tool, and final segment length definition that correspond to the 10 distinct dataset-splits. We statistically assess these slopes via the non-parametric sign test, by applying a 95% confidence level (p-value < 0.05), to obtain an overall per-dataset and per-tool slope trend. For p-values below 0.05 and a majority of slopes being positive, we classify the trend as *Predominantly Positive* (or simply *positive* in the following) as this indicates a consistent increase in the final testing log-likelihoods. Conversely, if the majority of slopes are negative (and the p-value is below 0.05), we classify the trend as *Predominantly Negative* (or *negative*), as this implies a proclivity that the corresponding tool overfits on the specific dataset. If neither of these conditions is met, we classify the overall trend as non-significant. Overall, we systematically evaluate overfitting, that is, we investigate whether overfitting emerges as *the* statistically expected behavior when optimization is pushed to its limits. This question is particularly relevant (a) for RAxML-NG and IQ-TREE, which employ thorough heuristics, and (b) in a comparative context, that is, when comparing overfitting trends in thorough versus faster (RAxML-NG ES, FastTree) heuristics.

Figure 2 presents exemplary learning curves for the anecdotal dataset 15034_1 obtained from RAxML-NG based inferences. In addition, the Figure displays the regression lines fitted to the final *n* = 30%*N* points of the testing curves. Compared to the DNN learning curves shown in Figure 1b, which illustrate the crossentropy loss during the DNN training example, the log-likelihood curves (especially the testing ones) are inherently more “bumpy”. As we discuss below, this behavior is rather expected.

Overall, our null model (*H*_0_ in statistical testing termi-nology) relies on two assumptions: First, we expect the same behavior (final curve) for the 10 holdout experiments (splits). This is well justified for comparatively long MSAs and supported by our random site selection approach, which ensures that each split constitutes a statistically representative sample of the MSA. Hence, the signal-to-noise ratio should remain unaltered across replicates, leading to analogous final curve behavior. Second, this behavior corresponds to a flat final curve (zero slope) that is perturbed by independent and identically distributed (i.i.d.) noise that has a null expectation and a symmetric distribution. This describes and captures the “bumpy” nature of our testing curves (Figure 2). Along the final curves of the ten experiments, we expect (under *H*_0_) the slope to exhibit an equal probability of being positive or negative (i.e., the basis of the sign test, which can also accommodate zeros when the slope is almost flat). The sign test is based on the binomial distribution with a probability of 0.5 and 10 trials under our settings. Since the number of trials is low, only contrasted results will be significant (e.g., 8 negative slopes, 1 positive slope, and 1 flat slope corresponds to a left-sided p-value of ~2%), while other cases will not be significant (e.g., 7, 2 and 1, respectively, p-value of ~9%).

Our approach focuses on the global trend in the final segment of the testing curve, which can be predominantly positive, negative, or non-significant. We specifically do *not* target the optimal point of the testing curve. This is because learning curves in deep learning are typically smooth as they are based on continuous numerical functions. In contrast, log-likelihood testing curves are inherently non-smooth (“bumpy”) because topological moves on the discrete tree parameter induce major log-likelihood changes/jumps (Figure 2), and the optimal point itself is therefore of reduced value. Consequently, focusing on the optimal point and comparing it with the final point (i.e., the output tree, which has a negative or null difference with the optimum) may result in overestimating overfitting trends and hinder the detection of under-optimization trends that correspond to a positive slope. This is, in fact, the main limitation we perceive in de Vienne et al. (2017). Nevertheless, we do include a comparison between the optimal and the final trees in the Discussion section (see Figure 7).

For simulated MSAs, we also generate topological accuracy curves by computing the quartet distances (Estabrook et al., 1985) between each intermediate sampled topology (*T*_*i*_), and the true tree topology (*T*_ref_). We use the quartet distance, rather than the more standard Robinson-Foulds distance (RF; Robinson and Foulds (1981)), because it is more refined, that is, it generates more possible, distinct values, and can therefore better capture fine-grained trends. We compute quartet distances (*d*(*T*_*i*_, *T*_ref_)) using the tqDist tool (Sand et al., 2014), and define the topological accuracy of each intermediate topology as 1− *d*(*T*_*i*_, *T*_ref_). We statistically evaluate the corresponding curves via the same frame-work as above (linear regression followed by the sign test). This analysis, however, does not directly assess overfitting. Topologies that diverge from the true tree can indeed yield equal, or better, log-likelihood scores on the testing sites while not degrading the model-fit. Analogous observations were reported in our previous works (Morel et al., 2020; Haag et al., 2022; Höhler et al., 2022; Togkousidis et al., 2023). While not focusing on cross-validation, we examined and confirmed the statistical plausibility of highly divergent topologies. The results suggest that ML tree inference on difficult-to-analyze MSAs as indicated by high Pythia scores, might yield topologically highly divergent, yet statistically equivalent topologies w.r.t. their likelihood score. The analysis of topological accuracy curves in the present paper provides novel insights into whether the inferred trees systematically converge toward (or diverge from) the true topology during the optimization process.

### 2.3. Overfitting Results

Figure 3 summarizes the results of the overfitting evaluation pipeline across 9,062 empirical (8,229 DNA and 833 AA) and 5,020 GTR+Γ-generated simulated DNA MSAs. For empirical MSAs, the proportion of negative and positive slope trends, as discussed in the following text, is visualized in Figure 4. We rarely observe systematic overfitting, as indicated by the proportion of negative slope trends.

**Figure 3.**
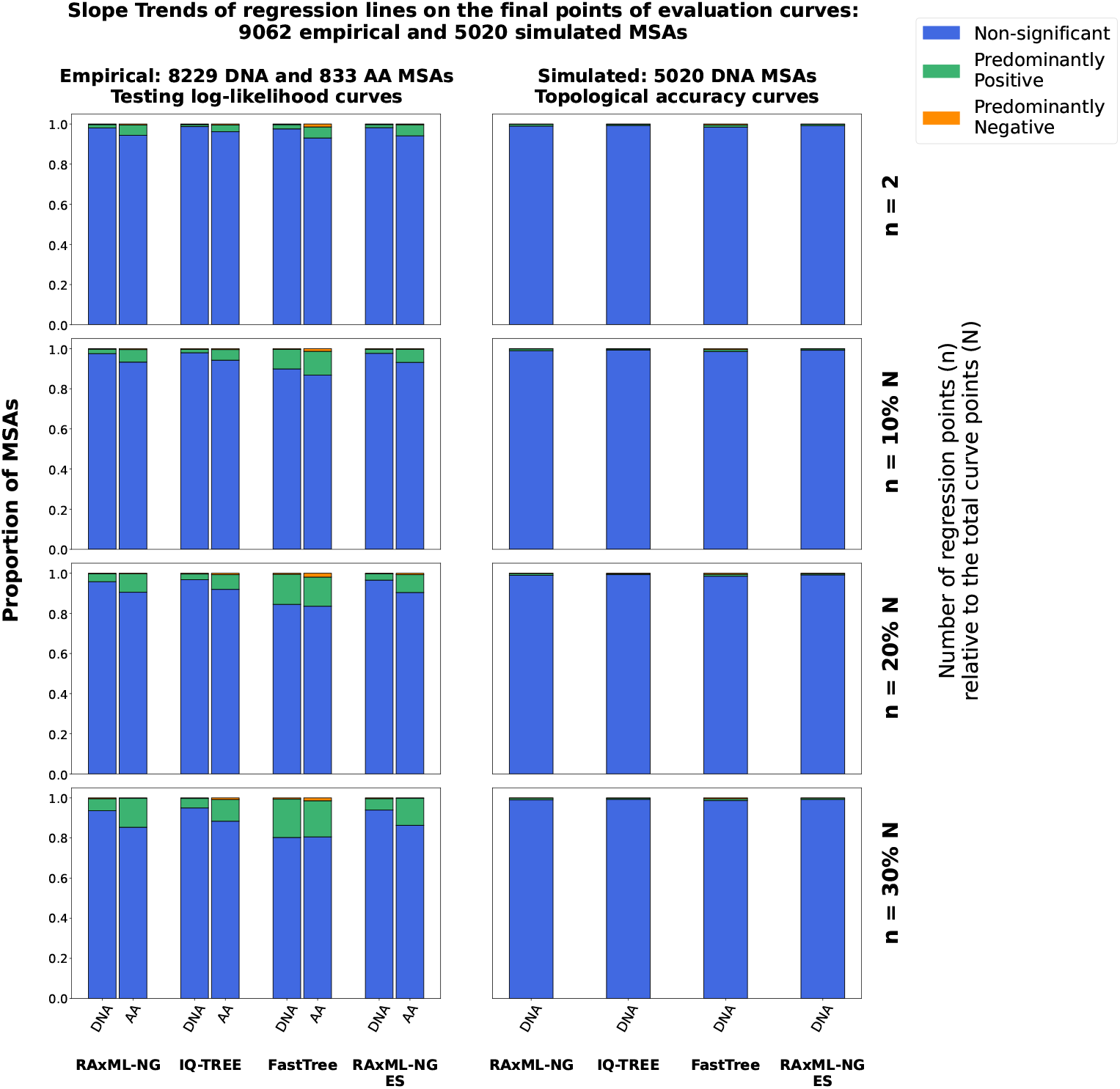
Slope trends of regression lines fitted to the final segments of evaluation curves for empirical (left) and simulated (right) MSAs. “Evaluation curves” refers to testing log-likelihood curves for empirical MSAs (left), and topological accuracy curves for simulated MSAs (right). Distinct columns within each subplot correspond to different ML tree inference tools: RAxML-NG, IQ-TREE, FastTree, and RAxML-NG ES. Subplot-rows correspond to different linear regression window sizes: the last 2 points, or the final 10%, 20%, or 30% of the points in each curve. The bar heights in the stacked bar charts indicate the proportion of MSAs exhibiting one of the three slope trends: Predominantly Positive, Predominantly Negative, or Non-significant, as classified via the non-parametric sign test.

**Figure 4.**
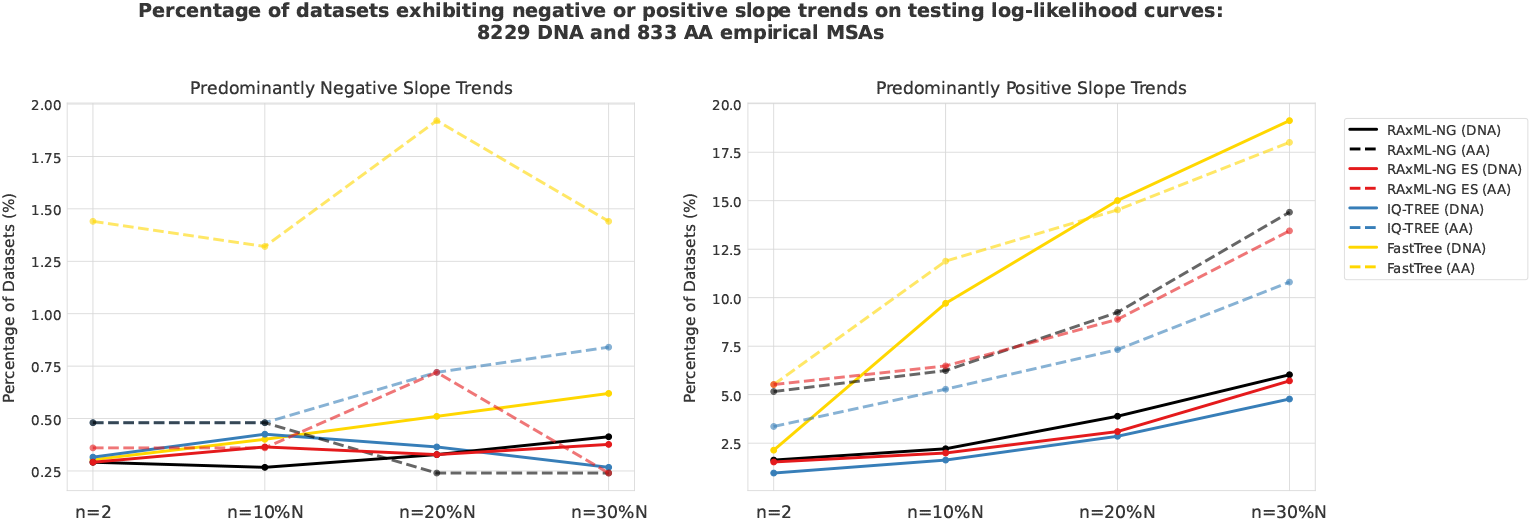
Percentage of datasets exhibiting Predominantly Negative (left) or Predominantly Positive (right) slope trends across different linear regression window sizes: the final 2 points (*n* = 2), the last 10% (*n* = 10%*N*), 20% (*n* = 20%*N*), or 30% (*n* = 30%*N*) of the points in each curve. The plot illustrates how the frequency of these slope trends varies across tools and regression window sizes, as those are observed in Figure 3, for empirical MSAs.

Focusing on the testing log-likelihood curves for empirical MSAs (Figure 3, left subplot), RAxML-NG, IQ-TREE, and RAxML-NG ES show highly consistent behavior across different regression window sizes. On DNA MSAs, we observe negative trends for approximately 0.3% of the MSAs, with only minor variations across tools and regression window sizes. For AA MSAs, the proportion remains similar (~ 0.3%) in most cases, except for IQ-TREE, where it increases to ~0.7%-0.8% for *n* = 20%*N* and *n* = 30%*N*. RAxML-NG ES also shows a slight increase to ~0.7% for *n* = 20%*N*. It subsequently drops to baseline levels for *n* = 30%*N*.

The above tools exhibit similar patterns for positive trends as well. Standard RAxML-NG and ES show positive trends on ~2% of DNA and ~5% of AA MSAs when only considering the final two points in the regression. As expected, the proportion of positive trends increases (roughly linearly) with the regression window size: to around 2.5% and 6% (DNA and AA MSAs) for *n* = 10%*N*, to 3.5% and 9% for *n* = 20%*N*, and to 6% and 14% for *n* = 30%*N*. IQ-TREE consistently yields lower positive trend rates: 1% and 3.3% (DNA and AA MSAs) for *n* = 2, 1.6% and 5.3% for *n* = 10%*N*, 3% and 7.3% for *n* = 20%*N*, and 5% and 11% for *n* = 30%*N*. Consequently, the proportion of non-significant trends ranges from ~94% and ~86% (DNA and AA MSAs, respectively) for *n* = 30%*N*, to ~98% and ~95% for *n* = 2. As expected, the proportion of non-significant trends increases when focusing on the very final parts of the curve that are closer to convergence, resulting in a corresponding positive trends decrease.

Note that the similarity between RAxML-NG and ES is expected. Early Stopping in RAxML-NG operates at the level of SPR rounds (see Methods) by omitting potentially redundant/unnecessary SPR rounds towards the end of the search and substantially reducing runtime. However, most SPR moves are accepted during the initial rounds. Therefore, in both RAxML-NG versions, a comparable number of intermediate topologies (i.e., curve points) is sampled and we only observe a modest reduction in intermediate topologies in RAxML-NG ES (see Supplementary Figure S10).

FastTree exhibits similar negative trends for DNA MSAs that range between 0.3% and 0.6%. For AA MSAs, however, the overfitting trends are substantially higher, with the baseline levels at approximately 1.4%, and increasing up to 1.9% for *n* = 20%*N*. These substantially higher negative trends for AA MSAs are difficult to interpret. They could be attributed to the plethora of approximations FastTree applies to calculate the log-likelihood scores of visited topologies, during the tree search (see Methods). These approximations can potentially misguide the search towards topologies that, when evaluated thoroughly on unseen sites, result in progressively decreasing testing log-likelihoods, for every intermediate accepted topology. Thorough tools, on the other hand, employ more accurate log-likelihood evaluations, which results in lower overfitting rates for AA MSAs. With respect to continuous parameter optimizations, RAxML-NG ES belongs to the group of thorough tools, since it invokes the same optimization routines as RAxML-NG, without applying any shortcuts (see Methods). Nevertheless, despite being inflated, the overfitting rates of FastTree are still small.

Regarding positive trends, FastTree exhibits relatively high rates for *n* = 30%*N* : 19.1% for DNA and 18% for AA MSAs. These rates gradually decline as we consider narrower final segments, reaching approximately 15% for *n* = 20%*N*, and 10%-12% for *n* = 10%*N*. This pattern suggests that slope trends remain strongly positive in the later stages of the tree search and indicates that FastTree could potentially benefit from additional topological optimization steps. When focusing on the final two points of the curve, though, FastTree attains levels that are comparable to the other tools, with 2.1% for DNA and 5.5% for AA MSAs. Hence, the proportion of non-significant trends in FastTree ranges from ~80% in both DNA and AA MSAs at *n* = 30%*N*, to 97.6% and 93% for *n* = 2 in DNA and AA MSAs, respectively. The observed convergence indicated by the decline of positive trends as we consider narrower regression segments, which is consistent among tools, does not imply that the quality of the inferred trees is equivalent. For instance, one way to assess the quality of inferred trees would be to compare them via statistical significance tests, as conducted in other studies (Höhler et al., 2022; Togkousidis et al., 2023, 2025). Such comparisons represent a static assessment of the model-fit observed in the output ML trees. In contrast, our study investigates dynamic overfitting trends, by independently analyzing the testing curves of each tool and evaluating whether model degradation is observed due to excessive optimization.

Additional results on empirical MSAs are presented in the Supplementary Material. First, in Supplementary Figure S4, we further evaluate the derived slope trends using the one-sample t-test, and the results are presented in juxtaposition to those obtained via the sign test. The outcomes are consistent across both tests and we do not observe any substantial differences. Second, for the specific case of *n* = 20%*N*, we investigate potential correlations between overfitting trends and MSA characteristics, such as the number of taxa, sites, patterns, and Pythia scores. The results are presented in Supplementary Figures S5–S7. A modest correlation suggests that overfitting trends are more prevalent for MSAs with relatively short sequences and low site pattern counts (fewer than ~10,000 sites and ~5,000 patterns), that is, as expected, with weaker phylogenetic signal. This upper sequence-length limit is particularly strict for RAxML-NG (both standard and ES versions), i.e., we observe no negative trends on MSAs exceeding those site/pattern thresholds. Overall, no clear correlation between higher difficulty scores and overfitting trends was observed (Supplementary Figure S7).

Regarding the 5,020 GTR+Γ-generated simulated DNA MSAs, we show the results of the topological accuracy curves, when evaluated using the same framework (linear regression followed by the sign test), on the right subplot of Figure 3. The overall trend is non-significant, exceeding 99% of the MSAs for RAxML-NG (standard and ES versions) and IQ-TREE, across all regression segment configurations. In FastTree, the proportion of non-significant trends is slightly lower, ranging between 98% and 99%, with the respective reduction being primarily added into negative trends. Specifically, while negative trends are observed in approximately 0.1% of the cases in RAxML-NG and IQ-TREE, FastTree consistently exhibits negative trends in 0.5% of cases, across all regression segment window sizes.

Additional results on simulated MSAs are presented in Supplementary Figures S8 and S9. The former illustrates the slope trends derived from the testing log-likelihood curves, and the latter from the topological accuracy curves. In both Figures, the results from the 5,020 GTR+Γ-generated datasets are presented in juxtaposition to those from the 1,322 JC-based simulated MSAs. As a reminder, we analyzed both types of simulated MSAs under GTR+Γ model in the ML tree inferences and evaluations (GTR+CAT for FastTree; see Methods) to also assess the effect of over-parametrization on JC-based MSAs. For the testing log-likelihood slope trends (Supplementary Figure S8), overfitting is essentially absent in both, the GTR+Γ- and JC-generated datasets across all tools (exact percentages are not reported, since they are negligible). JC-based MSAs display lower proportions of positive trends, primarily because these datasets tend to converge faster (as discussed in the next paragraph). Specifically, JC-based MSAs exhibit non-significant trends in ~99% of the cases, while the remaining 1% are attributed to positive trends. The slope trends derived from topological accuracy curves (Supplementary Figure S9) are consistent across both types of simulated data. Notably, the excess of free parameters induced by using the GTR+Γ model on JC-based MSAs does not increase overfitting trends.

Finally, the increased ratio of non-significant trends on simulated datasets can be explained by the fact that ML tree inferences generally converge more rapidly on such MSAs (Trost et al., 2024). The number of sampled topologies during ML inferences is substantially lower for simulated MSAs than for empirical ones, with the effect being particularly pronounced on JC-based MSAs. For instance, RAxML-NG visits on average twice as many intermediate topologies on empirical MSAs compared to GTR+Γ-simulated MSAs, and seven times as many compared to JC-simulated MSAs (see Supplementary Figure S10). There exist cases where the parsimony starting tree is identical to the final ML tree, that is, no topological moves are applied at all. In such cases, only one intermediate topology is sampled, and we set the corresponding slope to 0, biasing the slope data in favor of the non-significant outcome in the sign test. In this context, non-significant trends reflect easy convergence.

### 2.4. Holdout Validation (HV) in RAxML-NG

RAxML-NG HV inherently splits the input MSA into *model-training* (80%) and *validation* (20%) sites that are selected uniformly at random. Next, it constructs a parsimony starting tree, and conducts an ML tree inference exclusively on the model-training part of the MSA, while utilizing the validation sites to assess convergence and eventually terminate. More specifically, HV computes the log-likelihood scores on the validation MSA for the currently best topology after each SPR round (as well as on the initial parsimony tree). It terminates when the validating log-likelihood either decreases, or when the improvement is negligible (based on an *ϵ*-threshold of 0.1), for *k* successive steps. Upon termination, the topology with the highest *validating* log-likelihood is retrieved and printed to file. This is analogous to approaches used in deep learning, where a validation set is used to terminate the optimization process and prevent overfitting (Montavon et al., 2012). In the following, we execute HV for *k* := 1 and *k* := 5, and refer to the corresponding runs as HV-1 and HV-5.

Our experimental setup also follows the setup of a typical deep learning benchmarking experiment (Montavon et al., 2012), where the full dataset is partitioned into training and testing sets. The RAxML-NG versions under comparison (see below) use the training set for optimization, and the corresponding performance is evaluated on the testing set. The HV versions further account for overfitting, and they, in turn, split the training set into a model-training and a validation set, using the latter for convergence monitoring.

We benchmark RAxML-NG HV on 535 empirical (430 DNA and 105 AA) and 241 simulated (DNA only) long MSAs. We described the selection criteria for those MSAs at the beginning of the Results section. For each sampled MSA, we re-use the same 10-fold split we created for investigating topological overfitting, in the initial part of this work. Within each MSA-split, the RAxML-NG versions under comparison solely conduct inferences on the training MSA (using parsimony starting trees). The testing MSA exclusively serves as reference dataset for post-hoc evaluation. Specifically, for each individual MSA-split, we execute the following: (a) one standard RAxML-NG inference on the training MSA; (b) one ES inference on the training MSA; (c) one HV-1 inference, which further splits the training MSA into model-training (80%) and validation (20%) sites (the resulting MSAs comprise 64% and 16% of the original MSA sites, respectively); (d) one HV-5 inference, employing the same training MSA-partition as HV-1; and (e) one standard RAxML-NG inference on the model-training MSA (64%). We provide a rationale for this specific setup further below. We evaluate the 5 inferred ML trees, within each MSA-split, by applying the approximately unbiased (AU) test (Shimodaira, 2002), as implemented in the CONSEL tool (Shimodaira and Hasegawa, 2001), on the testing MSA. We thereby assess the output model-fit on unseen data. We mark the trees with a p-value exceeding 0.05 as being *plausible*. We aggregate the number of plausible ML trees inferred by each version across all splits, and record the number (out of 10) of plausible ML trees inferred by each version. We use this number as a metric to assess the overall performance across MSAs, that is, the higher the number of plausible ML trees inferred, the more robust the tool version is.

With the HV benchmark we address two main questions. The first is whether HV effectively evades overfitting by only capturing the phylogenetic signal of the input (training) MSA, while avoiding fitting inherent noise. Intuitively, a thorough inference (standard RAxML-NG) on the training MSA may fit both, signal and noise, mitigating the generalizability of the inferred trees on the unseen (testing) sites. Further, this effect should *a fortiori* be more pronounced when we further restrain standard RAxML-NG on the model-training MSA, which comprises only 64% of the original MSA sites. If this hypothesis is correct, HV is expected to yield trees with higher log-likelihood scores on the testing data, since it explicitly retains the tree with the highest validating log-likelihood, and thus maximizes the generalizability of the inferred model. Accordingly, we expect a fraction of HV trees to be statistically better compared to those inferred via standard RAxML-NG (on the training and model-training MSAs), when evaluated on the testing sites with the AU test.

The second question is whether HV provides a reliable strategy for Early Stopping in ML tree inference. To address this, we evaluate the relative performance of HV against that of ES, using standard RAxML-NG as the reference in both cases. The comparison considers both plausibility performance and runtime improvements.

We also compute quartet distances using tqDist, between the ML trees inferred by each version and the reference tree. On simulated MSAs, the reference tree is the true tree used to generate the data. On empirical MSAs, we consider as reference the ML tree inferred via standard RAxML-NG on the training MSA of each MSA-split.

### 2.5. RAxML-NG HV Results

Figure 5 illustrates the plausibility assessment results on empirical MSAs. The figure shows the fraction of empirical datasets, for which each tool version inferred *n* (out of 10) plausible ML trees, where *n* represents specific counts or intervals. Overall, the HV versions exhibit the weakest performance among all versions. Specifically, standard RAxML-NG infers 10 (out of 10) plausible ML trees for ~59% of DNA MSAs, ES for ~53%, the HV versions for ~35%, and RAxML-NG on the model-training MSA for ~40%. The corresponding values for AA MSAs are 64% for standard RAxML-NG, 55% for ES, 44% for the HV versions, and 47% for RAxML-NG (model-training MSA), respectively. The average number of plausible ML trees inferred per DNA dataset is ~9.3 for standard RAxML-NG and ES, 8.5 for the HV versions, and 8.7 for RAxML-NG on model-training MSAs; for AA MSAs, the corresponding averages are 9.4 for standard RAxML-NG, 8.2 for ES, 7.8 for HV versions, and 8.9 for RAxML-NG on model-training MSAs. Further, standard RAxML-NG runs (both on training and model-training MSAs) always infer at least one plausible tree (100%) on both DNA and AA MSAs. The remaining versions (ES and HV) fail to infer any plausible tree on 1 DNA and 2 AA MSAs, respectively. The failing DNA MSA is the same MSA for all three versions; the HV versions fail on the same two AA MSAs, one of which is also shared with ES. As expected, we observe a similar pattern in the plausibility assessment on simulated DNA MSAs (Supplementary Figure S12), although the differences are less pronounced, since simulated MSAs generally converge more consistently (Trost et al., 2024). Figure 6 illustrates the average quartet distance distributions between the ML trees inferred by each version and the corresponding true tree, on simulated DNA MSAs. The pairwise quartet distances, calculated between each ML tree inferred by each version, and the reference tree topology, are averaged over the 10 ML trees that each tool version infers per MSA. All distributions exhibit a median of 0, indicating that the average quartet distance is 0 for the majority of datasets across all methods. The mean values of the average quartet distance distributions are 0.014 for standard RAxML-NG and ES, and 0.017 for all other versions. Although HV versions and RAxML-NG (on model-training MSAs) exhibit identical mean values, the distribution of the latter appears to be slightly skewed towards zero values on the log-scale. This indicates that thorough RAxML-NG inferences that are constrained to the model-training sites only, almost outperform HV with respect to topological accuracy, even on such long simulated MSAs. Interestingly, a similar observation applies to ES relative to standard RAxML-NG (on training MSAs), with ES exhibiting a slight advantage. The performance difference between HV and the remaining versions becomes more pronounced on empirical MSAs (Supplementary Figure S13).

**Figure 5.**
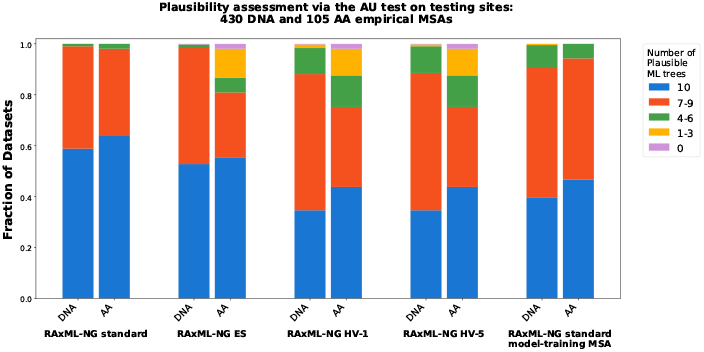
Plausibility test results on ML trees inferred via RAxML-NG versions on the training MSAs within each MSA-split, on empirical MSAs. We conduct this evaluation on the testing MSA of the corresponding MSA-split. We apply the AU test, as implemented in CONSEL, to the five ML trees inferred by: (a) RAxML-NG standard, (b) RAxML-NG ES, (c) HV-1, (d) HV-5, and (e) RAxML-NG standard on the model-training MSA of HV versions. ML trees with a p-value ≥ 0.05 are considered as being plausible. We aggregate the plausible ML trees across splits, and for each method and dataset, record the number (out of 10) of plausible ML trees.

**Figure 6.**
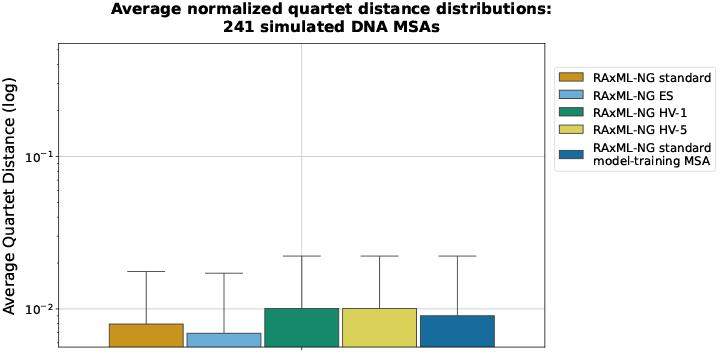
Average quartet distance distributions on simulated MSAs. We compute the quartet distance between ML trees inferred by each RAxML-NG version, and the true tree topology, on 241 simulated DNA MSAs. The y-axis is shown on a logarithmic scale.

Supplementary Figure S14 shows the speedup distributions of ES and HV relative to standard RAxML-NG. On both empirical and simulated MSAs, ES yields consistently better speedups than HV. These results provide a straightforward answer to the two questions posed above. First, HV does not offer any advantage regarding the prevention of overfitting. This aligns with our earlier findings that overfitting is insubstantial in ML-based phylogenetic inference. The use of a deep learning-inspired validation set thus appears inappropriate. Notably, standard RAxML-NG restrained to the model-training sites performs better than HV, despite using the same data fraction but without validation-based termination. Second, optimizing on all available sites yields better overall results, and runtime acceleration is more efficiently achieved via ES, which applies stopping rules that do not discard any data. On empirical DNA MSAs, ES performs analogously to standard RAxML-NG while yielding an average speedup of 3.5 ×. On AA MSAs, however, ES tends to terminate prematurely and underperforms with respect to standard RAxML-NG. This is consistent with our previous findings (Togkousidis et al., 2025). Our benchmark also includes two MSAs for which ES can not recover any plausible tree. Overall, our findings strongly suggest that holdout validation does not constitute a useful strategy in ML based phylogenetics, not even so on apparently sufficiently long MSAs.

## 3. Discussion

In the first part of our analysis, we investigated whether overfitting constitutes the statistically expected behavior of ML tree inference tools. We used the sign test to evaluate the statistical significance of the global trend among the final points of the testing curves. We evaluated RAxML-NG and IQ-TREE, which employ thorough heuristics, alongside with FastTree, a more superficial tool, and RAxML-NG ES, a recently introduced version of RAxML-NG that implements early stopping of the optimization process via the KH test combined with correction for multiple testing. Our findings show that, for empirical MSAs, overfitting, as indicated by negative slope trends, is only detected for less than 1% of the MSAs, across all tools and MSA types (DNA or AA). The only exception is FastTree on AA MSAs, where overfitting is observed for approximately 1.5% of the datasets. This might be attributed to the extensive approximations FastTree applies for log-likelihood calculations. Fast-Tree also exhibits an increased ratio of positive slope trends, suggesting premature termination of the optimization process before reaching a plateau. In such cases, further optimization might be beneficial.

Across all tools, the majority of empirical MSAs (above 80%) show non-significant slope trends. This pattern is even more pronounced on simulated MSAs, where non-significant trends dominate both, the testing log-likelihood, and the topological accuracy curves. In-terestingly, the excess of free model parameters introduced by analyzing the JC-generated simulated MSAs under the GTR+Γ model, does not induce topological overfitting. Instead, the tree inferences merely tend to converge more rapidly in such cases, potentially because the parsimony starting trees are more compatible with the JC model.

There may be instances, however, where our regression-based approach on the final parts of the testing curves fails to capture the actual model-fit degradation (overfitting), which manifests itself via a decrease in the testing log-likelihood score. One such scenario arises when the optimal point occurs too early in the testing curve, and is therefore excluded from the regression window (e.g., split 7 in Figure 2). Indeed, our sign test-based approach identifies strong tendencies specifically for the final points of the testing curves for each MSA. We deliberately did not compare the optimal point in the curve with the final one. This is because, due to the “bumpy” nature of the (testing) log-likelihood curves, the optimal point is of reduced value. Nevertheless, in Figure 7, we do present a two-point comparison between the corresponding optimal and the final points. The Figure comprises barplots that correspond to the accumulated number of positive, negative, and zero slope values, as observed across all 90,620 dataset-splits on empirical MSAs (10 for each MSA, across 9,062 MSAs), for *n* = 20%*N*. In addition to these slope type counts, the Figure also highlights the proportion of cases where the final ML tree is plausible (based on the AU test on the testing MSA) compared to the highest-scoring tree residing on the same testing log-likelihood curve. With this analysis, we statistically evaluate *significant* differences between the optimal and final trees, which constitutes a standard practice in ML tree comparisons (Goldman et al., 2000; Höhler et al., 2022; Togkousidis et al., 2023, 2025), rather than solely relying on likelihood differences. If the final tree is the highest-scoring one on the curve, we immediately consider it as being plausible. For RAxML-NG (both standard and ES), in 88% of dataset-splits the final tree is statistically indistinguishable from the optimal tree on the testing curve. In other words, under this plausibility test, overfitting is virtually absent in 88% of the cases. Moreover, in ~73% of dataset-splits with negative regression slopes (for *n* = 20%*N*), the final and the optimal trees are statistically indistinguishable. Interestingly, for dataset-splits with positive regression slopes, the final tree is statistically equivalent to the optimal tree in 95% of cases, implying that in the remaining 5% the final tree is significantly worse.

**Figure 7.**
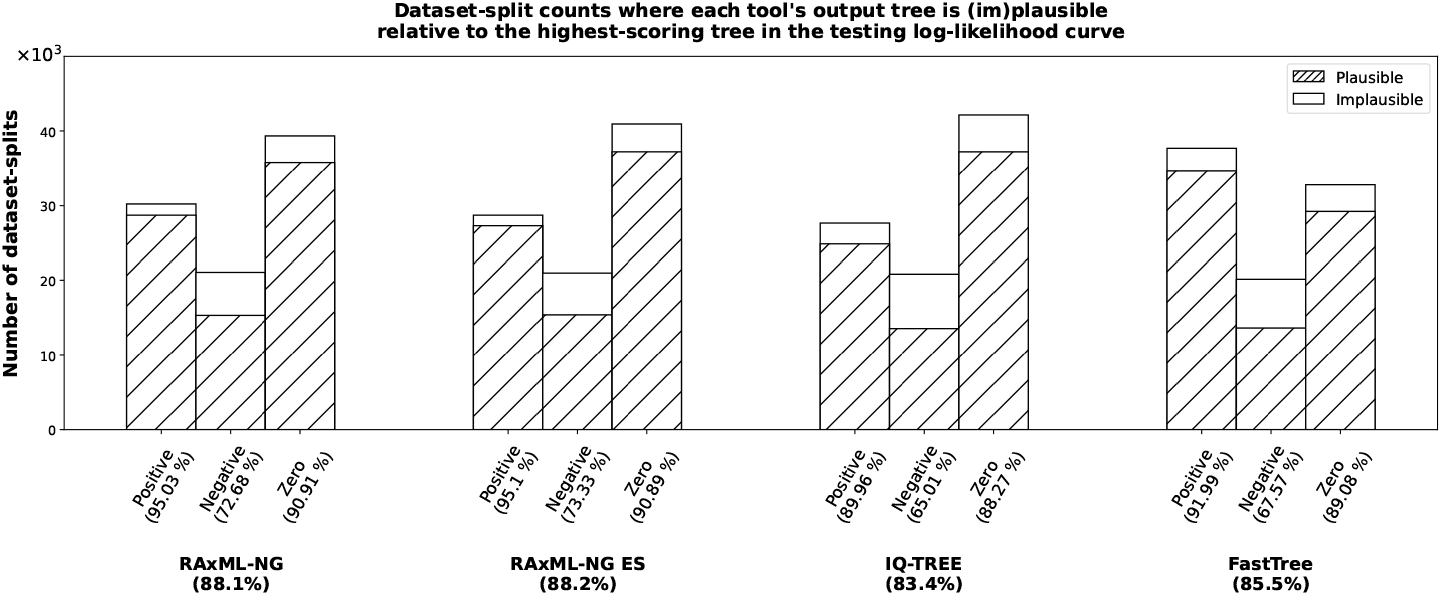
Barplot showing the accumulated number of positive, negative, and zero slope signs, across a total of 90,620 empirical dataset-splits. The slopes are derived from linear regressions on the final segments of the testing log-likelihood curves, for 9,062 empirical MSAs (10 splits per MSA). Bar heights represent the accumulated slope sign counts for *n* = 20%*N*. Results are shown separately for each phylogenetic inference tool. Within each bar, a distinct filling pattern indicates the percentage of cases where the final tree in the curve is plausible, compared to the highest-scoring tree (w.r.t. its log-likelihood on the testing MSA) on the same curve, as assessed via the AU test on the testing MSA. If the final tree is the highest-scoring of the curve, we directly mark it as plausible. We further report the percentages of plausible final trees in parentheses along the x-axis (for each slope sign category), and next to each tool label (aggregated across slope signs).

To explain instances where the final ML tree is non-plausible, despite of positive or zero slope signs, we consider two main factors: (a) strong oscillations along the curve (“bumpy” nature of the log-likelihood curves), and (b) scenarios where the highest-scoring tree occurs early on the curve and is excluded from the regression window (see Figure 2). These effects are, indeed, not captured by the sign test, and require additional trials as well as data points to be accurately assessed. This slight discrepancy, however, is expected when applying multiple statistical evaluation methods. On the other hand, the comparative approach depicted in Figure 7 has a specific limitation relative to the slope analysis using the sign test: because the optimal tree is always at least as good as the final tree, the two-point comparison can only yield two outcomes: (i) a significant difference, indicating overfitting, or (ii) a non-significant difference, indicating a likelihood-equivalence plateau. It is therefore unable to detect a perpetually increasing modelfit towards the final stages of optimization (i.e., positive slopes), which however is equally important when examining the dynamics of the testing curves, and thus tends to exaggerate overfitting trends.

At this point, we recap and synthesize our findings. Our sign-test analysis (Figure 3) showed that ML tools do not exhibit a strong tendency to overfit. Even when using thorough inference tools (RAxML-NG, IQ-TREE), topological overfitting is not statistically expected. At the same time, the two-point comparison (Figure 7) showed that in 12% of cross-validation splits (when using RAxML-NG), the model-fit degradation observed in the final tree is significant. This second result warns practitioners that, even if overfitting is not statistically expected in most cases, a downstream analysis of the inferred ML tree constitutes an important step of phylogenetic analysis. A fully resolved bifurcating tree can be over-parameterized, and some of its branches might not be supported by the data, as discussed in Berry et al. (2000). J. Felsenstein formalized this idea via the so-called *null-branch test* (Felsenstein, 2004). The null-branch test is a likelihood-ratio test (LRT) whose null hypothesis states that the length of a fixed inner branch is zero. If the null hypothesis is not rejected, the branch is collapsed and yields a polytomy. Building on this idea, the approximate likelihood-ratio test (aLRT; Anisimova and Gascuel (2006); Guindon et al. (2010b)) derives branch support values from likelihood comparisons and reports them as confidence-like measures on inner branches. An alternative approach is the standard bootstrap method (Felsenstein, 1985), which assesses the stability of inner branches under random resampling of alignment sites with replacement, and thus addresses a related, but conceptually distinct support notion. Several other methods have also been proposed to detect/eliminate overfitted branches (Mai and Mirarab, 2018; Sayyari and Mirarab, 2018; Shimodaira and Terada, 2019; Simmons and Kessenich, 2020; Zhang et al., 2021), while an alternative strategy introduces regularization via a penalized log-likelihood score (Kim and Sanderson, 2008). Overall, while the impact of topological overfitting is not expected to be large, practitioners should be aware of it and apply appropriate post-inference assessments.

Finally, our RAxML-NG HV benchmark highlights the limitations of holdout-based Early Stopping in ML tree inference. First, HV consistently underperformed in terms of both, statistical plausibility (based on the AU test), and topological accuracy, even when analyzing long MSAs. This is consistent with our previous observations, that is, that topological overfitting does not constitute an issue in ML tree inference. Second, the HV versions did not yield any run time speedups over the ES version. Therefore, randomly excluding a portion of the MSA during the inference, decreases the available phylogenetic signal and degrades the tree inference quality (Tan et al., 2015). We recommend practitioners to conduct ML inference using the full MSA, either via RAxML-NG for maximum accuracy, or the ES version as a faster, yet reliable alternative.

## 4. Materials and Methods

We initially present an overview of the heuristics employed by all ML inference tools we used. These tools are: RAxML-NG v1.2 (Kozlov et al., 2019), IQ-TREE v2.3 (Minh et al., 2020), FastTree v2.1 (Price et al., 2010), RAxML-NG Early Stopping (ES) (Togkousidis et al., 2025), and holdout validation (HV). All ML inference tools deploy heuristics, which are primarily organized into optimization blocks, comprising (a) topological optimization rounds, such as Subtree Prune and Regraft (SPR) or Nearest Neighbor Interchange (NNI) rounds, (b) Branch-Length Optimization (BLO), and (c) Model Parameter Optimization (MPO) rounds. We, further, describe the specific parameter configurations applied in the execution of each tool, and analyze the tool-dependent intermediate tree sampling approaches we developed, by detailing the necessary internal mechanics of their respective topological optimization rounds.

### 4.1. Tools

For the overfitting evaluation, we executed RAxML-NG and IQ-TREE under a thorough setting. Standard RAxML-NG uses a fixed log-likelihood improvement threshold (*ϵ*) to determine convergence of the tree inference and terminate. We set an *ϵ*-threshold of 0.1 (`--lh-epsilon 0.1` option), thereby protracting RAxML-NG’s tree search until the log-likelihood improvements between consecutive SPR rounds fall consistently below the specified value. In principle, when the log-likelihood improvement within an SPR round falls below the *ϵ*-value, RAxML-NG does not terminate immediately. Instead, it adjusts the SPR-specific parameters, either by increasing the minimum and maximum regrafting radii, or switching from FAST to SLOW rounds, following an MPO round. To approximate log-likelihoods of the visited topologies, FAST SPRs reuse existing branch lengths, whereas SLOW SPRs re-optimize the three branches adjacent to the insertion point. Full BLOs and MPOs are conducted on the initial and the final tree topologies. BLOs are also being executed after each SPR round, whereas only an intermediate MPO round is conducted between FAST and SLOW SPR rounds. Termination of the tree search occurs only after several consecutive SLOW rounds with varying SPR radii failed to yield any further improvement, relative to the specified *ϵ*-threshold (Kozlov, 2018).

An *ϵ*-value of 0.1 is considered a rather thorough setting in RAxML-NG. This value was the default in v1.1, yet was relaxed to 10 in v1.2, owing to the results of a systematic study (Haag et al., 2023), which showed that the original value was unnecessarily strict. Similarly, IQ-TREE uses a fixed *ϵ*-threshold of 0.1 to accept improvements in the current best-scoring topology; this parameter is not user-configurable. IQ-TREE only terminates after 100 consecutive NNI rounds fail to yield improvements, rendering its heuristic inherently exhaustive. Regarding continuous parameters, IQ-TREE performs full BLO on each “candidate tree” (see the following section for details), while thorough MPOs are conducted periodically, similar to RAxML-NG. We should further mention that, in all executions of RAxML-NG (including the ES and HV versions) and IQ-TREE, we accounted for rate heterogeneity across sites using four discrete gamma categories (Yang, 1994), thus applying the GTR+Γ model for DNA, and the LG+Γ model for AA MSAs. Overall, both tools employ thorough heuristics, combining strict *ϵ*-thresholds, exhaustive topological searches, and repeated optimizations of continuous parameters.

Investigating systematic overfitting under thorough heuristics can be informative with respect to whether the prolonged and compute-intensive optimization is justified, given the risk of inferring models that capture both signal and noise. To provide a contrasting perspective, we included FastTree in our comparison, which is a more superficial, “approximately-maximum-likelihood” Price et al. (2010) alternative. FastTree applies multiple shortcuts in both, discrete topological, and continuous parameter optimizations. During the first phase, the algorithm conducts minimum-evolution (ME; Rzhetsky and Nei (1993))-based NNI and SPR rounds, similar to FastME (Desper and Gascuel, 2002; Lefort et al., 2015), albeit using profiles instead of true distances to further accelerate computation. Additional heuristics confine the SPR search space from *O*(*m*^2^) to *O*(*m*) (where *m* is the number of taxa), and the total number of ME-based topological optimization rounds to 4 log_2_(*m*) NNIs, and at most 2 SPRs, respectively. In the second phase, FastTree performs ML-based NNI rounds that entail shortcuts in likelihood computations and confinements in NNI-space. FastTree also limits the total number of ML-NNI rounds to 2 log_2_(*m*). Partial BLOs under ML are conducted sporadically, while only a single MPO is conducted, between the ME- and ML-based topological optimization rounds. Furthermore, FastTree does not account for gamma-distributed rate heterogeneity during tree search. Instead, it employs a variant of the CAT approximation (Price, n.d.), which assigns each site to a fixed, precomputed rate category (from up to 20 categories). We say “a variant”, because FastTree’s CAT approximation differs from the original implementation in Stamatakis (2006). Further, the algorithm offers an option to optimize gamma rates (`-gamma` option), yet this is only applied to the output tree topology, and not during the topological optimization rounds. We disabled this option. We additionally reduced the number of fixed rate categories to 4 (`-cat 4` option) to intentionally further simplify the model approximations. Therefore, we employed the GTR+CAT4 model for DNA, and the LG+CAT4 model for AA MSAs. Overall, Fast-Tree serves as a fast, yet superficial alternative, which we opposed to thorough heuristics, to directly compare overfitting trends under both ML inference strategies. The final tool included in the overfitting assessment is RAxML-NG ES. Having already described the standard RAxML-NG heuristic, the exposition of the ES heuristic is straightforward. RAxML-NG ES performs a sequence of FAST SPR rounds followed by a sequence of SLOW SPR rounds; in both cases, the maximum subtree regrafting radius is fixed to 10. BLOs and MPOs are executed as in standard RAxML-NG. The key difference in ES, is the dynamic determination of the *ϵ*-threshold via the Kishino-Hasegawa (KH) test (Kishino and Hasegawa, 1989), which is applied to the two trees before and after each SPR round. The extracted p-value is corrected for multiple testing (`--stooping-criterion KH-mult` option) to account for the total number of SPR topologies that improve the likelihood relative to the best tree prior to the SPR round. When the log-likelihood improvement falls below this dynamically determined threshold, the algorithm either switches from FAST to SLOW SPR rounds, or terminates. Overall, ES provides a balance between thorough versus superficial heuristics, employing cautiously simplified topological optimization heuristics, while retaining full continuous parameter optimizations.

Finally, RAxML-NG HV is an iterative, holdout-based Early Stopping algorithm, analogous to holdout methods in deep learning. After parsing the input MSA, HV randomly splits the alignment into model-training and validation sites based on a user-defined ratio. The algorithm conducts tree inference on the model-training sites, and monitors convergence with the validation sites. HV first optimizes branch lengths and model parameters on the parsimony starting tree. Next, it conducts a series of FAST SPR rounds, evaluating each resulting topology on the validation MSA. HV interrupts this FAST-SPR sequence either when the validating log-likelihood drops, or when improvements fall below an *ϵ*-threshold of 0.1. In either case, HV retains the topology with the highest validating log-likelihood. The algorithm then conducts MPO and continues with a sequence of SLOW SPR rounds. Here, it terminates either when the validating log-likelihood decreases, or does not improve, for at most *k* consecutive rounds. We say at most *k*, because if no SPR move is performed within an SPR round, this indicates that the topology is already SPR-optimal on the model-training MSA, and repeating the SPR round is redundant. In such a case, HV interrupts the SLOW-SPR sequence, even before reaching *k* rounds. After the SLOW SPRs, HV retrieves and prints the topology with the highest validating log-likelihood to file. Further details regarding the exact command lines to invoke each tool and version are provided in the Supplement.

### 4.2. Tree sampling

Our initial strategy was to apply the testing sites, and thereby devise testing curves, on checkpoint trees printed by each tool under standard execution settings. However, differences in heuristics and in the frequencies of recording checkpoints resulted in incomparable testing curves. To resolve this, we reduced the tree sampling process to the most atomic level by storing *all* intermediate, accepted topologies which improve upon the current best log-likelihood score, as computed (i.e., approximated) by each tool. This sampling strategy ensures consistency across tools and improves curve resolution for overfitting assessment. However, the ML tools used in this study do not provide built-in options to record all accepted topologies in the manner we just described. Therefore, we modified the source code of each tool to implement our tree sampling strategy.

Implementing this fine-grained approach in RAxML-NG and FastTree was straightforward, whereas it proved more challenging for IQ-TREE. Both, RAxML-NG, and FastTree operate on a single tree during the tree inference, upon which they apply topological moves using greedy, hill-climbing heuristics. Within an SPR round in RAxML-NG, the algorithm prunes each subtree from the current tree and evaluates its possible re-insertions into neighboring branches, up to a certain radius from the pruning edge. The move that yields the highest score improvement, if such a move exists, is accepted; at this point we sample the updated topology, and the algorithm proceeds to the next subtree. The same sampling strategy is adopted in RAxML-NG ES, since the two versions invoke the same SPR round function, differing only w.r.t. to the input parameters. FastTree follows an analogous greedy approach but applies additional shortcuts. First, it performs ME-SPR moves, accepting only moves that improve the ME score. We sampled all such accepted topologies. Next, it conducts both ME- and ML-based NNI moves, during which the algorithm compares the scores of the three NNI-neighboring topologies around a central, inner branch, accepting the topology with the highest score and moving to the next inner branch; we sample every accepted topology.

IQ-TREE exclusively uses NNI moves to navigate through tree space. During an NNI round, IQ-TREE visits and evaluates all NNI-neighboring topologies of the current tree, and sorts them based on their score improvements. Starting from the highest-ranked move, the algorithm sequentially applies all *compatible* NNI moves to the current tree. A set of NNI moves is compatible, if the central branches between any two moves in the set are non-adjacent. When applying those compatible NNIs, the log-likelihoods of intermediate occurring topologies are not computed; only the final topology is properly evaluated including BLO. If the final likelihood exceeds the score of the highest-ranked NNI move on the initial list, the resulting topology is accepted; otherwise, the applied NNI moves are reversed, except for the first move, which guarantees the highest score improvement. This process is repeated iteratively, until no further NNI moves yield score improvements. This heuristic was initially introduced in Guindon and Gascuel (2003), and is referred to as the hill-climbing NNI round in the original IQ-TREE paper (Nguyen et al., 2014).

NNI-based heuristics, however, are particularly prone to become trapped in local optima. To overcome this, IQ-TREE employs two additional mechanisms; the candidate tree set (CTS) and the perturbation step. The CTS is a fixed-size population of locally (sub-)optimal trees maintained by the algorithm. At the beginning of every round, the algorithm randomly draws a tree from the CTS and applies random NNI moves to it (perturbation), to escape from the local optimum. The perturbed tree serves as input for the next hill-climbing NNI round. After each NNI round, both, the CTS, and the best-found ML tree are updated accordingly. The process is repeated and the algorithm terminates only after the best-found topology has remained unchanged for a predefined number of consecutive rounds (100 by default).

Given the additional complexity in the IQ-TREE heuristic, it becomes challenging to reduce the tree sampling process to its most atomic level. Specifically, in cases where the log-likelihood of the resulting topology after the application of multiple, sequential, compatible NNI moves improves upon the score of the currently best-found ML tree, it is unclear at which move this so far best log-likelihood was surpassed. Sampling only the final topology, which is guaranteed to improve the log-likelihood, may result in undersampling. Consequently, the sampling strategy that we followed for IQ-TREE involves sampling *all* intermediate topologies encountered during the sequential application of compatible NNI moves, when this set of moves yields an update in the best-found ML tree. This strategy might introduce a minor oscillation in the training curve. While the resulting training curves may resemble a stochastic optimization process, the generic trend remains increasing, capturing even marginal score improvements resulting from topological moves.

Finally, the current study investigates overfitting trends for each tool independently. That is, the devised testing curves are evaluated separately for each tool. The consistent tree sampling strategy among tools allows for meaningful cross-comparisons and enables us to examine the effect of the specific heuristics employed by each method. The tool-dependent approximations applied in the log-likelihood score calculations of each visited topology, directly affect the search trajectories. For instance, RAxML-NG and IQ-TREE rely upon more accurate likelihood calculations than FastTree, thereby guiding the search towards topologies that do not systematically degrade model-fit, whereas FastTree exhibits such a degradation more frequently. However, the testing curves devised by RAxML-NG and FastTree exhibit greater similarity than those from IQ-TREE, rendering them directly comparable. This is because the devised testing curves comprise an incremental sequence of SPR-neighboring topologies, where each accepted topology is an SPR-neighbor of its predecessor, yielding a smooth tree space trajectory. This is not the case for IQ-TREE, and the apparent discontinuities in IQ-TREE testing curves may complicate direct comparisons between IQ-TREE and the other tools.

## Data and Software Availability

We implement both RAxML-NG HV, and the tree sampling algorithm, in RAxML-NG v1.2. These modifications are available as open source code under GNU GPL at https://github.com/togkousa/raxml-ng/tree/overfitting-early-stopping. This repository further includes the modified version of FastTree v2.1 that includes the sampling algorithm. FastTree is added as a separate folder in the same repository and is available as open source code under the GNU GPL. Finally, our sampling algorithm has been integrated into IQ-TREE v2.3 and is available as open source code under the GNU GPL at https://github.com/togkousa/iqtree2/tree/overfitting. Invocations for all tools are detailed in the Supplementary Material. All datasets and results generated in this study are publicly available at https://cme.h-its.org/exelixis/material/overfitting_data.tar.gz.

## Acknowledgment

We thank Damien M. de Vienne for his constructive feedback, which helped us to improve the overall quality of the manuscript. This study was financially supported by the Klaus Tschira Foundation and by the European Union (EU) under Grant Agreement No 101087081 (Comp-Biodiv-GR). O.G. was supported by PRAIRIE (ANR-19-P3IA-0001).

