## Supplementary Material for "A Systematic Investigation of Overfitting in Maximum Likelihood Phylogenetic Inference"

[1] Computational Molecular Evolution Group, Heidelberg Institute for Theoretical Studies, Germany; [2] Institute of Theoretical Informatics, Karlsruhe Institute of Technology, Karlsruhe, Germany; [3] Institut de Systématique, Evolution, Biodiversité (ISYEB, UMR7205 - CNRS, Muséum National d'Histoire Naturelle, SU, EPHE, UA), Paris, France; [4] Biodiversity Computing Group, Institute of Computer Science, Foundation for Research and Technology, Greece

### Introduction

This Supplement provides additional information and results to complement the main text. First, we provide details regarding the datasets used, pipeline stages, command-line instructions for the invocation of all tools, and our HPC cluster configuration. Next, we present additional analyses and plots discussed in the main text, which further support and extend our findings.

### Datasets - Experimental pipeline

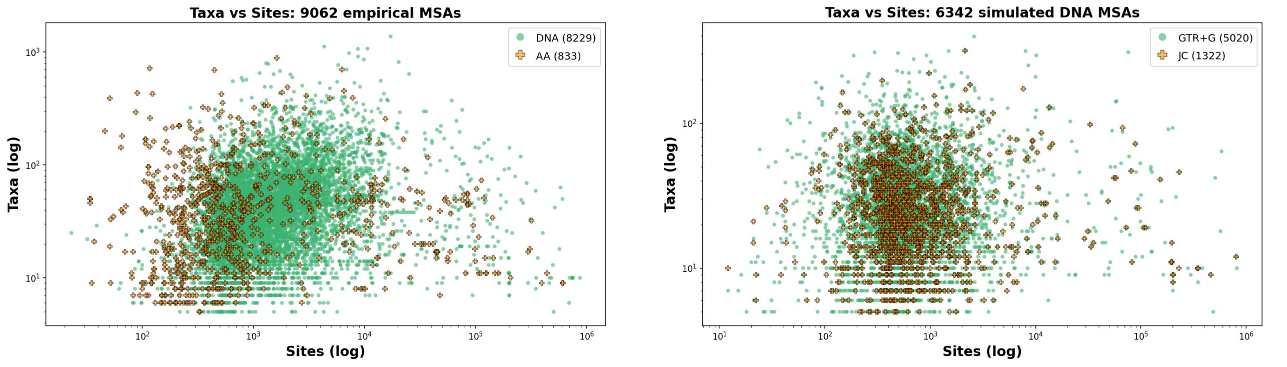


(a) (b)

**Figure S1.** Scatter plots showing the number of taxa and sites for 9,062 empirical and 6,342 simulated MSAs. **(a)** Empirical MSAs: 8,229 DNA (circles) and 833 AA (crosses) datasets. **(b)** Simulated MSAs: 5,020 generated under the GTR+$\Gamma$ model (circles), and 1,322 under the JC model (crosses).

As described in the main text, our data collection comprises 9,062 empirical MSAs (8,229 DNA and 833 AA) from TreeBASE (Piel et al., 2009), and 6,342 simulated DNA MSAs from a prior benchmark study (Trost et al., 2024). Regarding the simulated MSAs, 5,020 of these were generated under the GTR+$\Gamma$ substitution model, and 1,322 under the JC model, using the AliSim tool (Ly-Trong et al., 2022). Figure S1 illustrates the numerical properties (number of sites and taxa) of the entire collection of empirical and simulated MSAs.

Figure S2 shows density histograms of the Pythia score distributions for the 9,062 empirical and 6,342 simulated MSAs. The distributions are generally shifted toward lower Pythia scores, suggesting easier MSAs. We attribute this shift to the preprocessing that we applied to all datasets, as described in the main text. Specifically, we removed duplicate taxa and gap-only columns. This preprocessing and data cleaning step improved the overall phylogenetic signal strength. To highlight this effect, we mention two characteristic anecdotal cases of DNA MSAs: dataset 11756_1, whose Pythia score decreased from 0.62 to 0.13, after removing 36 duplicated sequences and 4 gap-only columns, and dataset 11269_0, whose Pythia score decreased from 0.84 to 0.04, after removing 22 duplicated sequences. For comparison, the original Pythia score distribution of the unprocessed TreeBASE MSAs is closer to uniform (see Figure 5 in Togkousidis et al. (2023)). Further, MSAs with difficulty scores above 0.8 are rare. This reflects an inherent limitation in the definition of the Pythia (difficulty) score, as previously discussed in the Supplementary Material of Togkousidis et al. (2023).

Figure S2 illustrates the Pythia score distribution for the subset of MSAs used to benchmark RAxML-NG HV, which corresponds to lighter-colored bars. As mentioned in the main text, these distributions are even further shifted toward lower difficulty scores, consistent with our intentional selection of long, statistically informative MSAs for holdout-validation benchmarking.

A schematic representation of the experimental pipeline used for overfitting evaluation is provided in Figure S3. It is important to note that preprocessing the MSAs was mandatory to ensure the smooth execution of all tools included

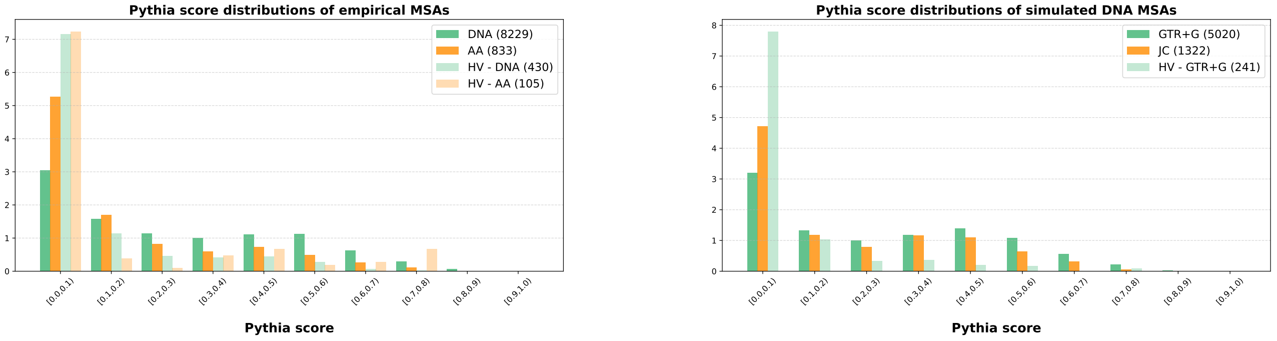


(a) (b)

**Figure S2.** Density histograms showing the Pythia score distributions for 9,062 empirical and 6,342 simulated MSAs, as well as for the 535 empirical and 241 simulated MSAs selected to benchmark RAxML-NG HV. Datasets sampled for holdout-validation benchmarking are depicted via lighter-colored bars. **(a)** Empirical MSAs: 8,229 DNA and 833 AA datasets, along with 430 DNA and 105 AA MSAs selected for RAxML-NG HV benchmarking. **(b)** Simulated MSAs: 5,020 generated under the GTR+$\Gamma$ model and 1,322 under the JC model. The 241 MSAs used for benchmarking were all generated under the GTR+$\Gamma$ model.

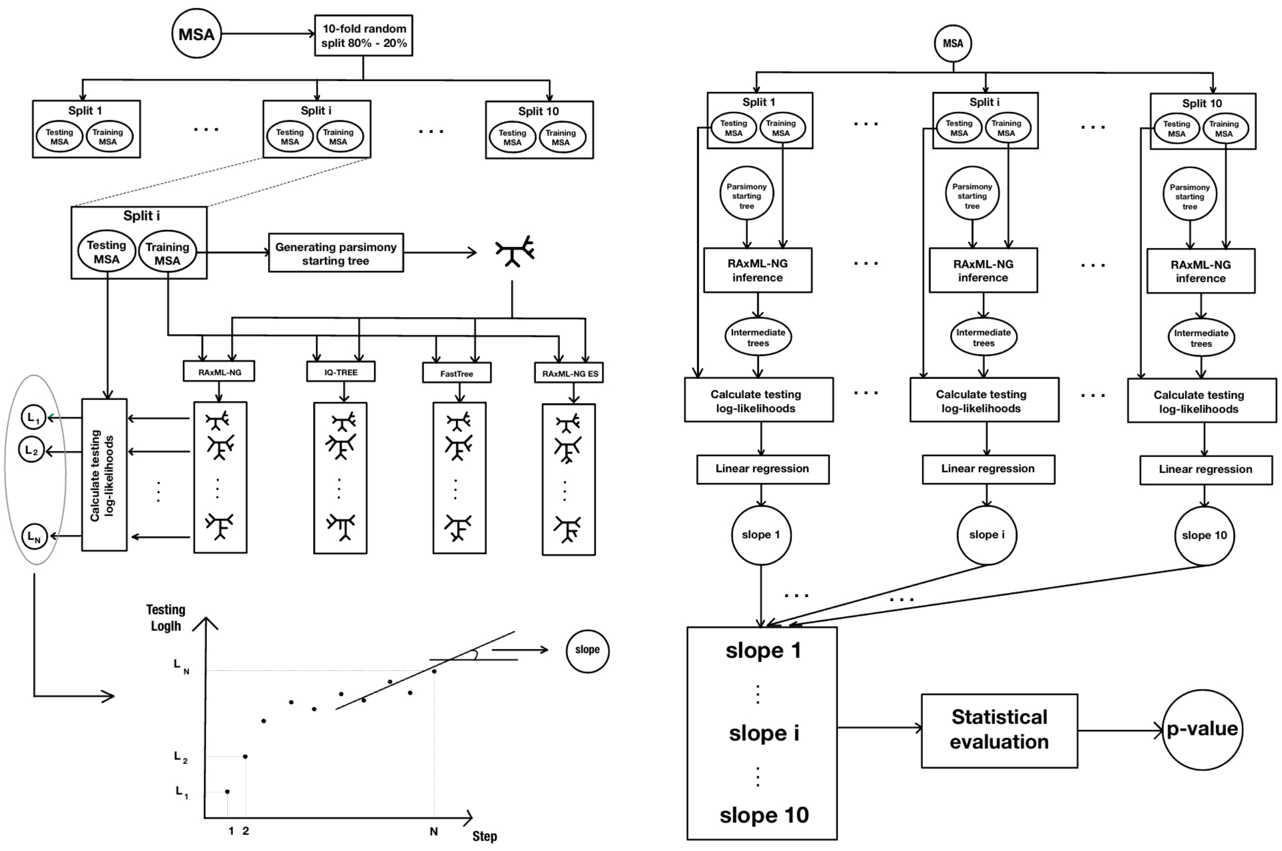


(a) (b)

**Figure S3.** A schematic representation of the overfitting evaluation pipeline. **(a)** Each MSA is randomly partitioned into training and testing sites, applying a 10-fold Monte-Carlo cross-validation approach, with a splitting ratio of 80%-20%. For each dataset-split, we use the training sites to generate a parsimony tree via RAxML-NG, which serves as the starting point for RAxML-NG, IQ-TREE, FastTree, and RAxML-NG ES to conduct ML tree inferences on the training sites. During each individual ML tree inference, we sample the intermediate improved (w.r.t. the log-likelihood score) topologies, accepted by each tool. Next, we use the testing MSA to compute the testing log-likelihoods of the sampled topologies in order to construct testing curves. We perform a linear regression on the final points of each testing curve and extract the slope of the fitted line. **(b)** A more abstract overview of the entire process. At the bottom, the 10 slopes of the regression lines (corresponding to the 10 datasets-splits) are collected and statistically assessed using the non-parametric sign test. The figure illustrates the process for RAxML-NG only, we apply it analogously to IQ-TREE, FastTree, and RAxML-NG ES.

in the pipeline. This is because the ML inference tools handle duplicated taxa and gap-only columns inconsistently. For example, we observed that IQ-TREE might fail to execute when gap-only columns are present. In addition, IQ-TREE, by default, automatically shrinks alignments to MSAs containing unique sequences and therefore samples topologies with fewer taxa than expected. This subsequently leads to errors when these trees are being evaluated with RAxML-NG on the comprehensive MSA. While FastTree generally tolerates duplicated rows (sequences), the sampled intermediate topologies for datasets comprising duplicates typically contain multifurcations. However, RAxML-NG fails to evaluate multifurcating trees, as it can only process strictly bifurcating trees, which causes the pipeline to terminate with an error. Thus, to avoid the need for tool-specific exception handling and ensure consistency across all analyses, we uniformly preprocessed all datasets by removing duplicated taxa and gap-only columns. Furthermore, during the random partitioning of MSAs into training and testing sets, we ensured that the training MSAs only contained unique taxa. If duplicated taxa occurred in the training subset due to random splitting, the partitioning was repeated until a valid split was obtained. For testing MSAs, we did not filter duplicates, as RAxML-NG effectively handles alignments with duplicated sequences, during the testing log-likelihood evaluations.

### Commands and bwForCluster Helix details

We implemented the tree sampling algorithm for RAxML-NG (both for the standard and the ES version), as well as the HV algorithm, in the `overfitting-early-stopping` branch of a forked RAxML-NG repository[1]. The same repository also contains the modified version of FastTree in a separate folder. The tree sampling modification of IQ-TREE is implemented in the `overfitting` branch of a distinct forked IQ-TREE repository[2].

#### RAxML-NG v1.2:

Tree sampling in RAxML-NG is enabled using the `--chkpt-method 2` option. In our experiments, we explicitly set the likelihood improvement thresholds via the options `--lh-epsilon 0.1 --lh-epsilon-triplet 1`. The models we used are GTR+$\Gamma$ (`--model GTR+G` option) for DNA MSAs, and LG+$\Gamma$ (`--model LG+G` option) for AA MSAs. Parallelization is implemented and tested for this version. Below is the command used to conduct one ML tree inference on a training MSA (e.g., `training.phy`) using 4 threads:

```
./raxml-ng-adaptive --adaptive off --threads auto{4} --msa training.phy --model {model}
--tree pars{1} --chkpt-method 2 --lh-epsilon 0.1 --lh-epsilon-triplet 1 --prefix standard
--seed 0
```

In the example command, the adaptive heuristic (Togkousidis et al., 2023) is disabled using the `--adaptive off` option. For convenience, we use the prefix `standard` in our command. RAxML-NG stores the intermediate improved topologies in the `standard.raxml.sprTrees` file, and the parsimony starting tree in `standard.raxml.startTree`. This starting tree is subsequently used as input for the remaining tools.

We, further, note that we used RAxML-NG v1.2 to evaluate intermediate trees sampled from all ML tools during the overfitting assessment, using the corresponding testing MSAs. As described in the main text, in this evaluation we re-optimized branch lengths and substitution model parameters. To slightly accelerate this evaluation, we applied an $\epsilon$-threshold of 1.0. Below we provide an example command to evaluate the stored intermediate improved topologies on a testing MSA (e.g., `testing.phy`), using 4 threads:

```
./raxml-ng-adaptive --evaluate --threads 4 --msa testing.phy --model {model}
--tree standard.raxml.sprTrees --lh-epsilon 1.0
```

#### RAxML-NG ES:

To invoke the KH-multiple testing stopping criterion, we use the command `--stopping-criterion KH-mult`. When using this command, the adaptive heuristic is automatically disabled. Further, in the ES version, we specify no $\epsilon$-threshold. Below is the command used to invoke RAxML-NG ES, using 4 threads and the prefix `es`:

```
./raxml-ng-adaptive --threads auto{4} --msa training.phy --model {model}
--tree standard.raxml.startTree --chkpt-method 2 --prefix es --seed 0
```

RAxML-NG ES stores the intermediate improved topologies in the `es.raxml.sprTrees` file.

[1]https://github.com/togkousa/raxml-ng/tree/overfitting-early-stopping
[2]https://github.com/togkousa/iqtree2/tree/overfitting

##### Modified IQ-TREE:

One can enable the tree sampling in our modified IQ-TREE version using the `-wt` flag. Parallelization also works for this modified IQ-TREE version. Below is the command used to invoke our modified IQ-TREE version, using 4 threads and the prefix `iqtree` (for example, on a DNA dataset):

```
./iqtree2 -nt 4 -s training.phy --seqtype DNA -m GTR+G -t standard.raxml.startTree
--prefix iqtree -wt --seed 0
```

The modified IQ-TREE version stores the intermediate improved topologies in the `iqtree.treels` file.

##### Modified FastTree:

The tree sampling is automatically enabled in our modified FastTree version. As we mentioned in the main text, FastTree uses a variant of the CAT approximation (Stamatakis, 2006; Price, n.d.) to model rate heterogeneity across sites by assigning each site to one of 20 fixed rate categories by default. To further simplify the model approximations, we reduced this number to 4, using the `-cat 4` option. Therefore, we used the GTR+CAT4 model for DNA (`-gtr -nt -cat 4` option), and the LG+CAT4 model for AA MSAs (`-lg -cat 4` option). Below is the command used to invoke the modified FastTree version on a DNA dataset:

```
./FastTree -gtr -nt -cat 4 -intree1 standard.raxml.startTree -log fasttree.log < training.phy
> fasttree.bestTree
```

In the above command, the output log file is set to `fasttree.log`. Modified FastTree uses the prefix of the logfile (here, `fasttree`) to name the sampled topologies file. In this example, our modified FastTree version stores the intermediate improved topologies in the `fasttree.chkpTrees` file.

##### RAxML-NG HV:

Users can invoke the holdout-validation algorithm in RAxML-NG via the `--holdout-es` flag. The algorithm randomly splits the input MSA into model-training and validation sites, based on the specified seed, and using a default splitting ratio for the model-training MSA of 0.8 (i.e., 80% of the sites). One can define an alternative splitting ratio via `--split-ratio X`, where `X` is a floating point number between 0 and 1 that specifies the proportion of *model-training* sites. Further, the user can also specify the number of convergence iterations $k$ (see main text) via `--conv-iters k`, where `k` must be a positive integer value. The default value of convergence iterations is $k := 1$. The resulting model-training and validation MSAs can be stored by enabling the `--extra split-save-phylip` option. The MSAs are written to the `{prefix}.raxml.training.phy` and `{prefix}.raxml.testing.phy` files, respectively. Parallelization is implemented and tested for this version. An example RAxML-NG HV invocation, where we use $k = 5$ convergence iterations (HV-5; see main text), is:

```
./raxml-ng-adaptive --holdout-es --threads auto{4} --msa {model} --model {model}
--conv-iters 5 --prefix example --extra split-save-phylip --seed 0
```

RAxML-NG HV stores the resulting model-training and validation MSA files in the `example.raxml.training.phy` and `example.raxml.testing.phy` files, respectively.

##### Computing Environment

We executed the overfitting evaluation pipeline on the bwForCluster Helix, located at the Heidelberg University Computing Centre. We utilized the `cpu-single` cluster partition, which allocates submitted jobs to AMD nodes. Each AMD node is equipped with two AMD Milan EPYC 7513 Processors, running at 2.60GHz, providing a total of 64 physical cores per node. The operating system is RedHat Linux. Further, we benchmarked RAxML-NG HV on one of our lab servers, equipped with two AMD EPYC 9684X (Genoa-X) processors, each with 96 physical cores and two threads per core, operating at a maximum frequency of 3.72 GHz. The operating system is Ubuntu 24.04.

#### Overfitting analysis: Supplementary results

Figures S4–S11 provide additional results supporting the overfitting assessment discussed in the main text. Figure S4 compares slope trend classifications on 9,062 empirical MSAs, based on two statistical tests: the non-parametric sign test (as used throughout the analysis presented in the main text) and the one-sample t-test. Figures S5–S7 illustrate the observed slope trends on 9,062 empirical MSAs for the case of $n = 20\% N$, against MSA characteristics, such as sites and taxa (Figure S5), sites and patterns (Figure S6), and the Pythia score (Figure S7). As mentioned in the main text, a modest correlation suggests that overfitting trends are more prevalent for MSAs with relatively short sequences and low site pattern counts (fewer than ∼10,000 sites and ∼5,000 patterns). This sequence-length limit is particularly strict for RAxML-NG. In contrast, for IQ-TREE and FastTree, a small number of DNA MSAs exhibiting overfitting trends were detected as outliers, despite their high sites-over-taxa and patterns-over-taxa ratios. One notable outlier, which was observed in FastTree, was the DNA MSA 13654_5. This dataset exhibited negative trends despite comprising 10 taxa, 747,575 sites, and 58,956 patterns that render its phylogenetic signal extremely strong (Pythia score 0.0). A direction for a future work would be to focus on MSAs that exhibit systematic overfitting and identify MSA-specific characteristics that predispose particular tools to overfit. Comparative analyses may also be informative, for example by detecting MSA features that trigger overfitting in RAxML-NG versus IQ-TREE. Based on the current results, the two tools do not overfit on the same MSAs. For $n = 20\%N$, RAxML-NG shows negative (overfitting) trends in 29 MSAs and IQ-TREE in 36, yet the overlap comprises only 4 MSAs. Three of these MSAs show strong phylogenetic signal (Pythia score below 0.1), and the fourth is still relatively easy to analyze (Pythia score 0.29).

Figures S8) and S9) extend the overfitting analysis to 6,342 simulated DNA MSAs. Figure S8) shows slope trend distributions for testing log-likelihood curves across 5,020 GTR+$\Gamma$- and 1,322 JC-generated DNA MSAs, while Figure S9) reports the same trends based on topological accuracy curves. Further, Figure S10 summarizes the distribution of the number of intermediate topologies sampled per tool and dataset, across all empirical and simulated MSAs. Exact numerical results from the above Figures are not reported here, as those have already been discussed in the main text, and any observed difference is either minor, or not sufficiently informative to necessitate a dedicated mention.

Figure S11 shows the distribution of normalized non-zero slope values across all 90,620 dataset-splits. Normalization is necessary because MSAs vary in length, and absolute log-likelihood differences are not directly comparable. Each slope value is divided by $0.2 \cdot s \cdot t$, where $s$ is the number of sites in the corresponding MSA, $t$ is the number of taxa, and the $0.2$ factor is appended because the testing log-likelihoods are computed on testing sites which account for 20% of the full alignment length. The result is multiplied by $10^5$, to express the values as log-likelihood differences per hundred thousand MSA characters (i.e., the number of sites multiplied by the number of taxa). Whisker lengths in boxplots are adjusted, such that ~10% of values are considered as outliers: about 4–4.5% are negative outliers (below the lower whisker), and 5.5–6% are positive outliers (above the upper whisker). The figure shows that, among ML inferences that yield non-zero slope values during the final segments of the testing curves, the normalized slopes range from approximately -100 to +100 slope units (i.e., log-likelihood difference per topology per $10^5$ MSA characters). In such cases, overfitting may be asserted from negative slopes, as they reflect model-fit degradation. However, based on the results presented in Figure 6 of the main text, the log-likelihood differences in 73% of the instances exhibiting negative slopes in RAxML-NG, are not statistically significant when assessed via the AU test.

Finally, Table S1 reports the accumulated number of positive, negative, and zero slope values, observed across all 90,620 dataset-splits of all 9,062 empirical MSAs (10 splits per MSA). The table also reports the average number of each slope type per dataset. Our results show that, across 10 inferences per MSA, an average number of 4 to 6 inferences result in a zero slope at the final segment of the curve. The only exception is FastTree that shows a majority of positive slopes for $n = 20\%N$ and $n = 30\%N$. This aligns with our earlier observation that FastTree might benefit from additional optimization. Further, positive slopes represent the second most frequent slope type, both in terms of accumulated counts, and average per-dataset values. On average, 0.5 to 1.5 more inferences per MSA show positive, rather than negative slope values.

#### Limitations in de Vienne *et al.* preprint

In the main text, we stated that de Vienne et al. (2017) was the first study to investigate topological overfitting as a dynamic phenomenon. Our study was inspired by this work. However, we believe that the preprint exhibits certain limitations. We briefly mention two of them. First, the authors evaluated overfitting trends in PhyML (Guindon et al., 2010) and classic RAxML (Stamatakis, 2014). To sample intermediate topologies, they used the `--print_trace` command in PhyML and the `-j` command in RAxML. While the PhyML command indeed stores all intermediate topologies in a way analogous to our procedure, the RAxML command records only a subset of intermediate topologies (i.e., those inferred after each SPR round). This important detail is documented in the RAxML 8 manual[3]. This means that (a) not every topology-induced improvement is captured for the RAxML curves, and (b) the resolution of the resulting curves differs between PhyML and RAxML which complicates a direct comparison.

Second, the authors selected the best tree (in terms of testing log-likelihood) among the final ones, which they compared to the very last tree, the former being inevitably better (or at least not worse). This introduces a strong bias towards negative slope trends, and may over-estimate the overfitting signal. In our procedure, we circumvent this potential bias. Therefore, our results differ from those reported by de Vienne *et al.*. Specifically, we decrease this potential bias by simply selecting the two last points ($n = 2$), or all last points within a given fraction ($n = 10\%$, $20\%$ or $30\%$).

[3]https://cme.h-its.org/exelixis/resource/download/NewManual.pdf

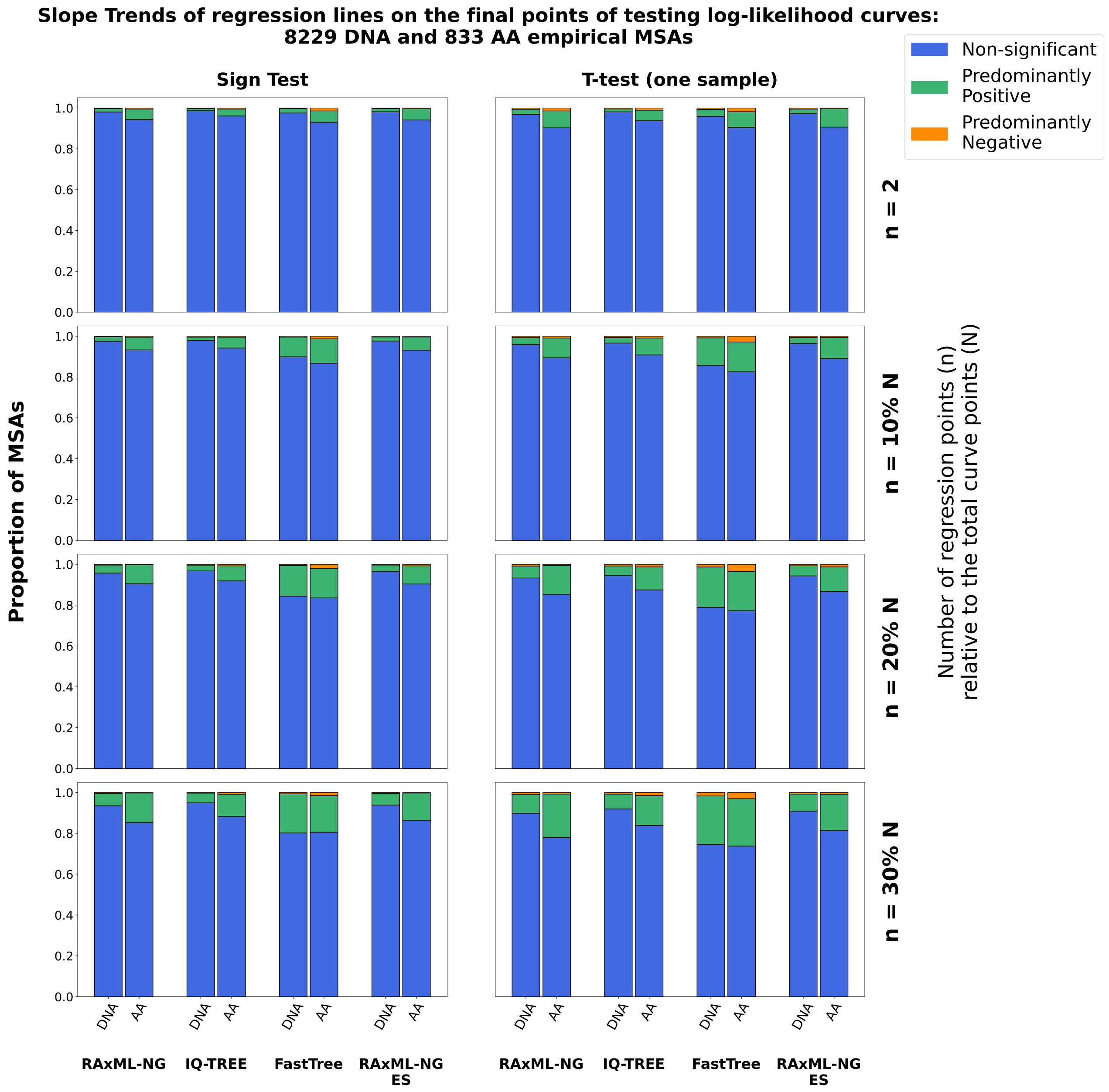


**Figure S4.** Slope trends of regression lines fitted to the final segments of the testing log-likelihood curves, derived from 9,062 empirical MSAs. The slope trends are evaluated using either the non-parametric sign test (left), or the one-sample T-test (right). The sign test results (left subplot) correspond to those presented in the main text (Figure 2, left subplot). Distinct columns within each subplot correspond to different ML tree inference tools: RAxML-NG, IQ-TREE, FastTree, and RAxML-NG ES. Subplot-rows correspond to different linear regression window sizes: the last 2 points, or the final 10%, 20%, or 30% of points in each curve. The bar heights in the stacked bar charts indicate the proportion of MSAs exhibiting one of the three slope trends: Predominantly Positive, Predominantly Negative, or Non-significant, as inferred by the statistical tests used.

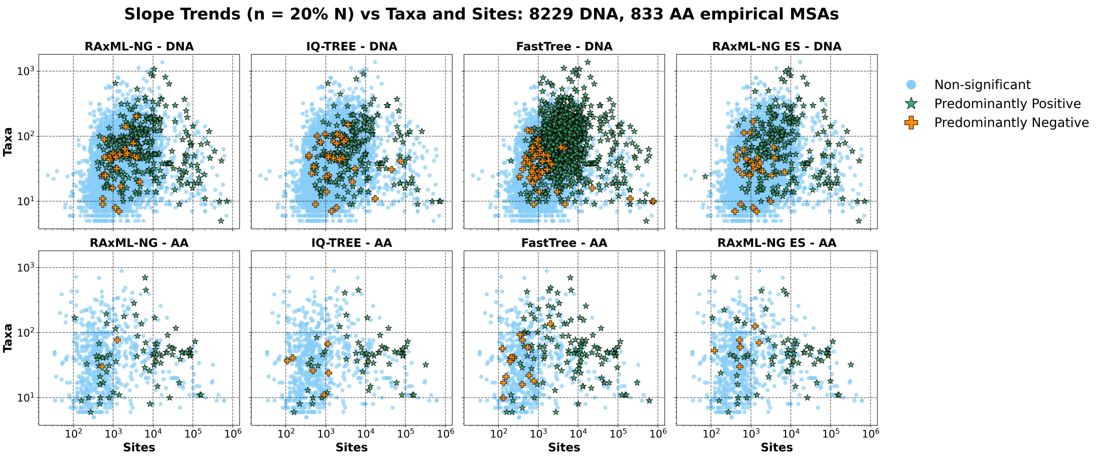


**Figure S5.** Slope trends versus taxa and sites in 9,062 empirical MSAs, illustrated for the specific case where $n = 20\%N$, with $n$ denoting the number of regression points and $N$ the total number of points on the testing log-likelihood curves. Both horizontal and vertical axes are shown on a logarithmic scale.

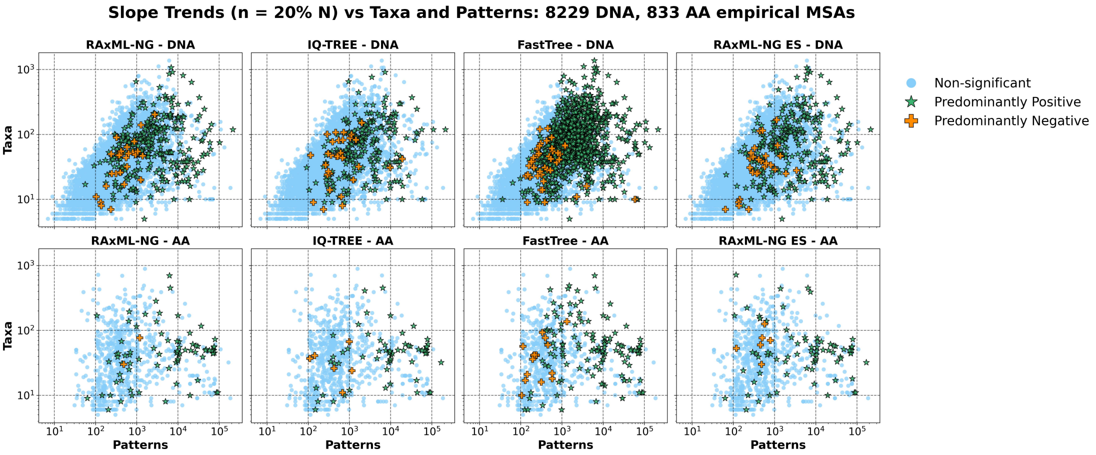


**Figure S6.** Slope trends versus taxa and patterns in 9,062 empirical MSAs, illustrated for the specific case where $n = 20\%N$, with $n$ denoting the number of regression points and $N$ the total number of points on the testing log-likelihood curves. Both horizontal and vertical axes are shown on a logarithmic scale.

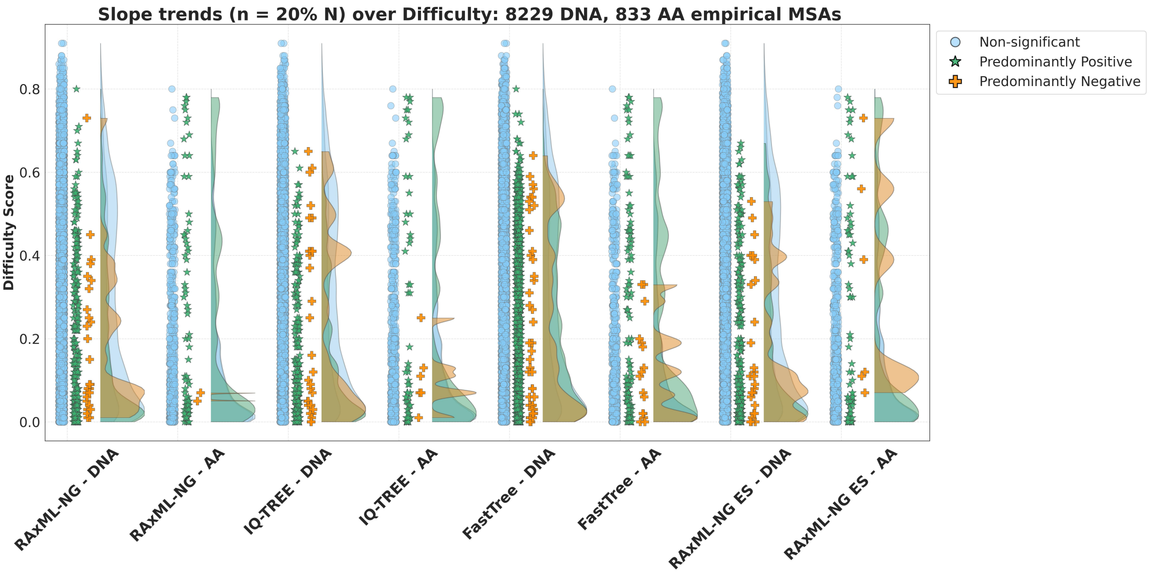


**Figure S7.** Slope trends over difficulty scores for 9,062 empirical MSAs, illustrated for the specific case where $n = 20\%N$, with $n$ denoting the number of regression points and $N$ the total number of points on the testing log-likelihood curves.

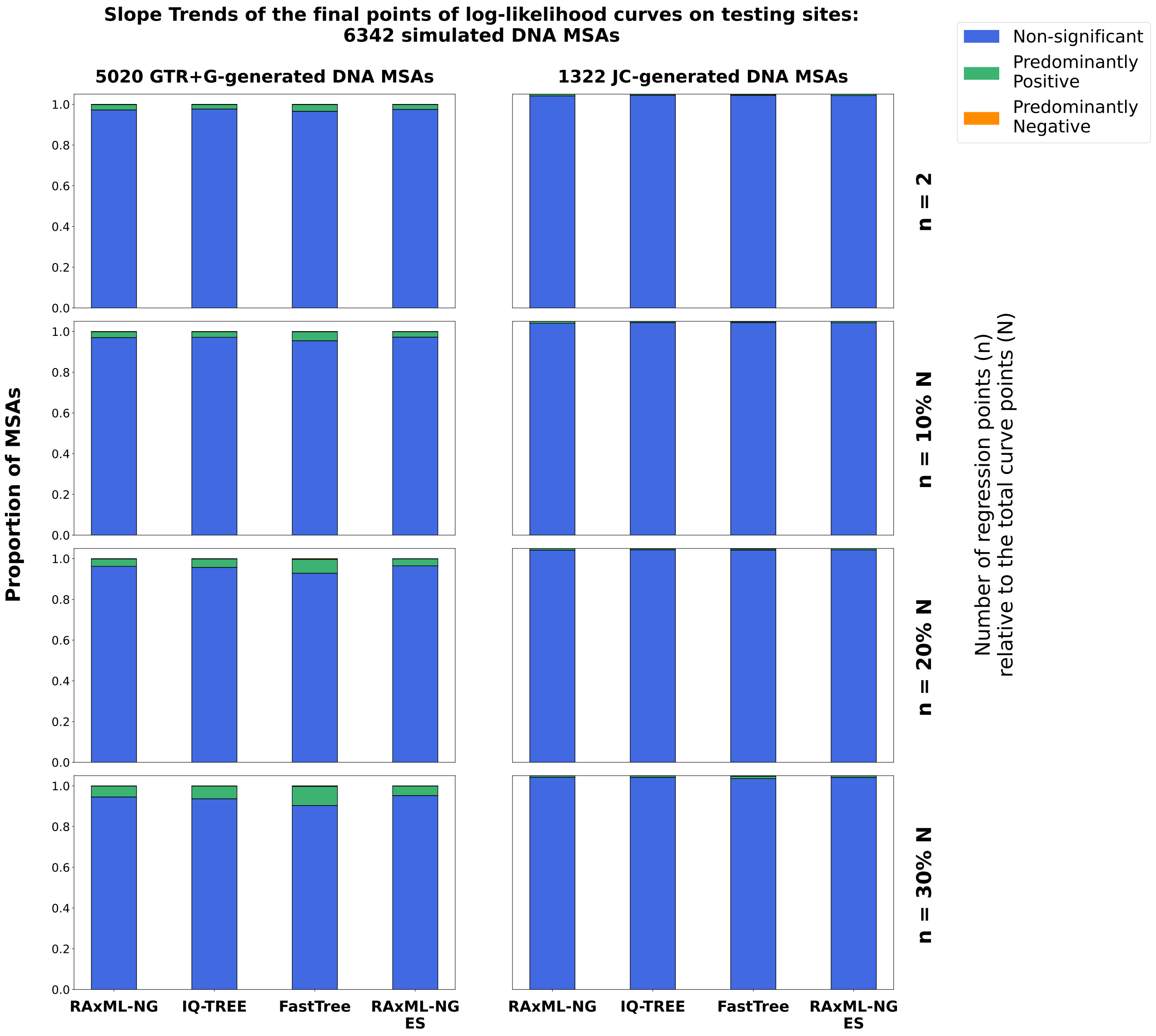


**Figure S8.** Slope trends of regression lines fitted to the final segments of the testing log-likelihood curves, derived from 6,342 simulated DNA MSAs, comprising 5,020 GTR+$\Gamma$-generated (left), and 1,322 JC-generated MSAs. Distinct columns within each subplot correspond to different ML tree inference tool: RAxML-NG, IQ-TREE, FastTree, and RAxML-NG ES. Subplot-rows correspond to different linear regression window sizes: the last 2 points, or the final 10%, 20%, or 30% of points in each curve. The bar heights in the stacked bar charts indicate the proportion of MSAs exhibiting one of the three slope trends: Predominantly Positive, Predominantly Negative, or Non-significant, as inferred by the non-parametric sign-test.

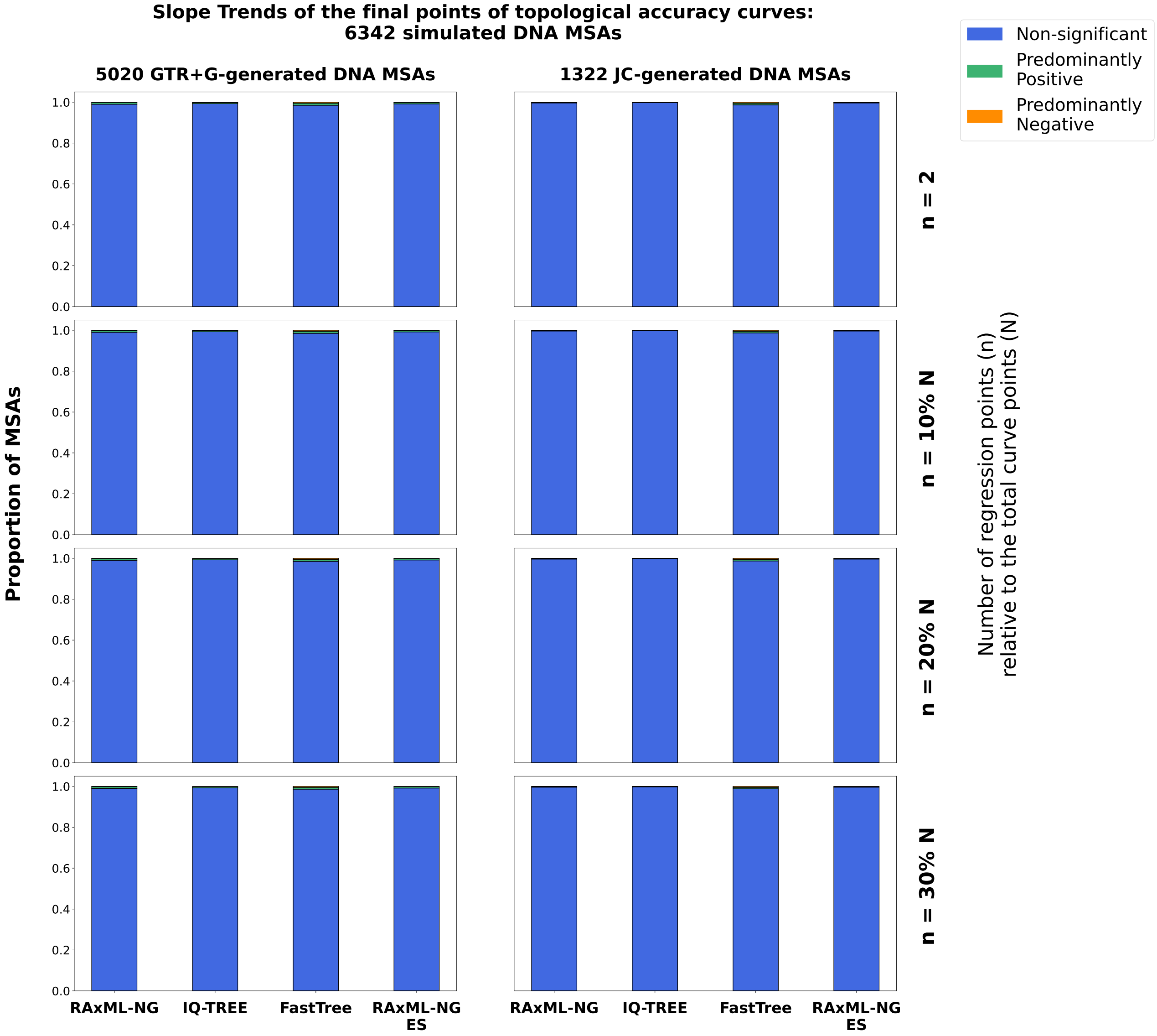


**Figure S9.** Slope trends of regression lines fitted to the final segments of the topological accuracy curves, derived from 6,342 simulated DNA MSAs, comprising 5,020 GTR+$\Gamma$-generated (left) and 1,322 JC-generated MSAs. Results of GTR+$\Gamma$-generated MSAs (left subplot) correspond to those presented in the main text (Figure 2, right subplot). Distinct columns within each subplot correspond to different ML tree inference tool: RAxML-NG, IQ-TREE, FastTree, and RAxML-NG ES. Subplot-rows correspond to different linear regression window sizes: the last 2 points, or the final 10%, 20%, or 30% of the points in each curve. The bar heights in the stacked bar charts indicate the proportion of MSAs exhibiting one of the three slope trends: Predominantly Positive, Predominantly Negative, or Non-significant, as inferred by the non-parametric sign-test.

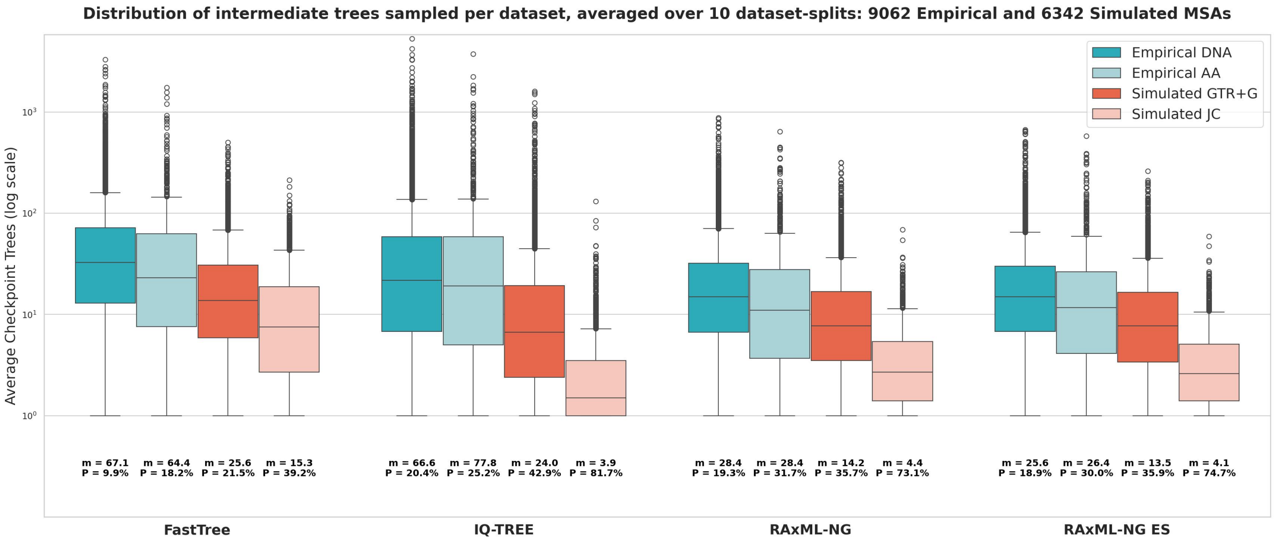


**Figure S10.** Distributions of the number of intermediate trees sampled by each tool for each dataset, averaged over the 10 dataset-splits, across 9,062 empirical and 6,342 simulated MSAs. For each boxplot, the mean value of the distribution, and the percentage (P) of MSAs with an average number of sampled topologies below 5 are reported. The y-axis is shown on a logarithmic scale.

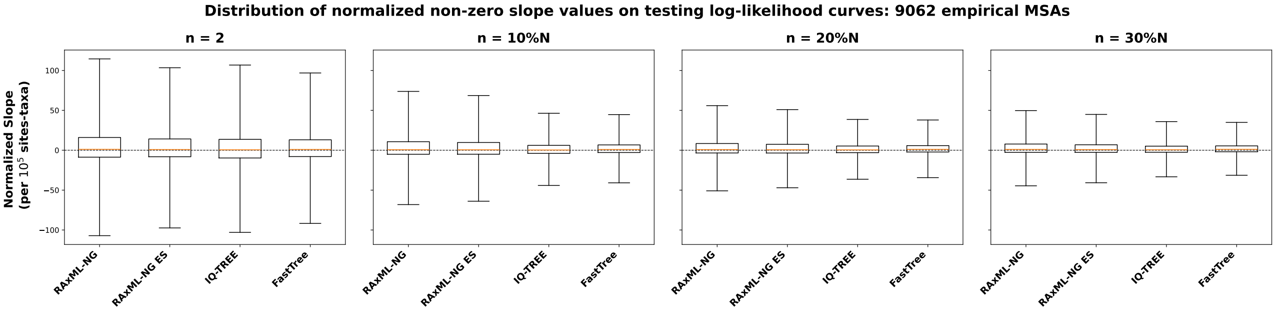


**Figure S11.** Distributions of normalized non-zero slope values on testing log-likelihood curves, derived from 90,620 dataset-splits across 9,062 empirical MSAs. Slope values are normalized to correspond to $10^5$ sites-taxa, ensuring comparability across MSAs of varying lengths.

| | n = 2 | | | n = 10%N | | | n = 20%N | | | n = 30%N | | |
|---|---|---|---|---|---|---|---|---|---|---|---|---|
| **Tool** | **Pos** | **Neg** | **Zero** | **Pos** | **Neg** | **Zero** | **Pos** | **Neg** | **Zero** | **Pos** | **Neg** | **Zero** |
| RAxML-NG | 22,519 / 2.48 | 17,827 / 1.97 | 50,274 / 5.55 | 26,029 / 2.87 | 20,042 / 2.21 | 44,549 / 4.92 | 30,230 / 3.34 | 21,052 / 2.32 | 39,338 / 4.34 | 33,720 / 3.72 | 21,173 / 2.34 | 35,727 / 3.94 |
| RAxML-NG ES | 21,648 / 2.39 | 17,640 / 1.95 | 51,332 / 5.66 | 24,697 / 2.72 | 19,429 / 2.14 | 46,494 / 5.13 | 28,725 / 3.17 | 20,959 / 2.31 | 40,936 / 4.52 | 32,550 / 3.59 | 21,200 / 2.34 | 36,870 / 4.07 |
| IQ-TREE | 16,597 / 1.83 | 14,500 / 1.60 | 59,523 / 6.57 | 24,096 / 2.66 | 19,828 / 2.19 | 46,696 / 5.15 | 27,671 / 3.05 | 20,804 / 2.29 | 42,145 / 4.65 | 30,230 / 3.33 | 20,359 / 2.25 | 40,031 / 4.42 |
| FastTree | 20,641 / 2.28 | 16,353 / 1.80 | 53,626 / 5.92 | 32,104 / 3.54 | 19,248 / 2.12 | 39,268 / 4.33 | 37,675 / 4.16 | 20,131 / 2.22 | 32,814 / 3.62 | 40,627 / 4.48 | 20,376 / 2.25 | 29,617 / 3.27 |

**Table S1. Accumulated number of positive ("Pos" columns), negative ("Neg"), and zero ("Zero") slope signs, across 90,620 empirical dataset-splits in total**, derived from linear regressions on the final segments of the testing log-likelihood curves, for 9,062 empirical MSAs (10 splits per MSA). The table further reports the average number of slopes being positive, negative, and zero per dataset. Each row corresponds to a distinct tool, and columns are grouped into four sections, each representing a distinct regression window: $n = 2$, $n = 10\%N$, $n = 20\%N$, and $n = 30\%N$. Each cell reports two values in the format X / Y: X is the accumulated number of regression lines exhibiting the specified slope sign, and Y is the corresponding average per dataset.

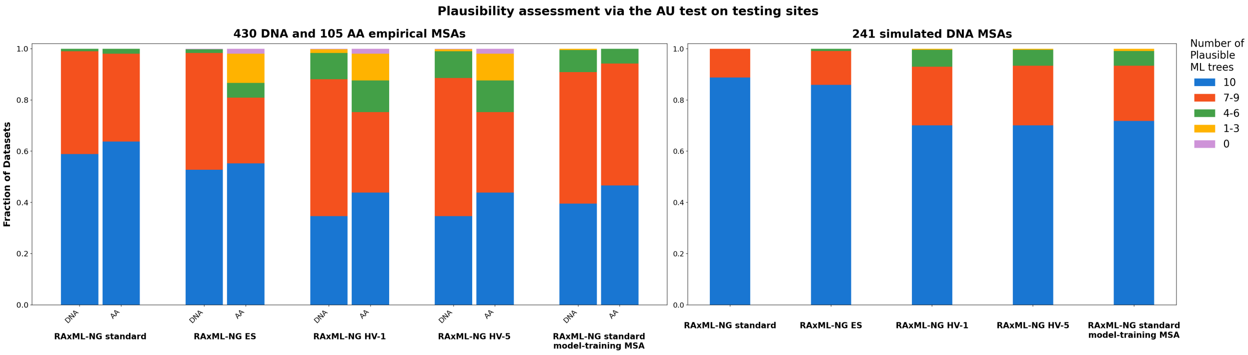


**Figure S12.** Plausibility test results on ML trees inferred via RAxML-NG versions on the training MSAs within each MSA-split, on empirical (left) and simulated (right) MSAs. The results on empirical MSAs (left subfigure) correspond to those already presented in the main text (Figure 4). We conduct this evaluation on the testing MSA of the corresponding MSA-split. We apply the AU test, as implemented in CONSEL, to the five ML trees inferred by: (a) RAxML-NG standard, (b) RAxML-NG ES, (c) HV-1, (d) HV-5, and (e) RAxML-NG standard on the model-training MSA of HV versions. ML trees with a p-value $\geq 0.05$ are considered as being plausible. We aggregate the plausible ML trees across splits, and for each method and dataset, record the number (out of 10) of plausible ML trees.

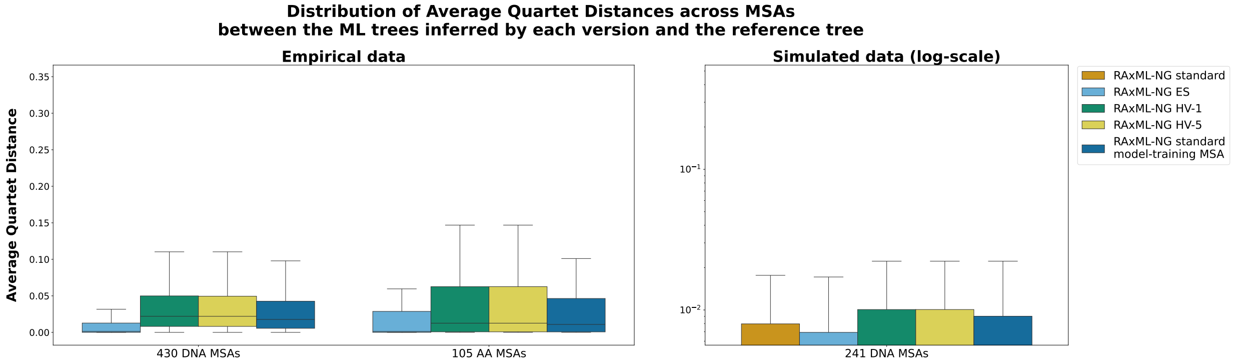


**Figure S13.** Average Quartet Distance distributions between the ML trees inferred by each RAxML-NG version, and the reference tree topology, for empirical (left subfigure) and simulated (right subfigure) MSAs. For empirical MSAs, the reference tree is the ML tree inferred by standard RAxML-NG on the training MSA of the corresponding MSA-split. Consequently, the distribution of standard RAxML-NG (on training MSAs) is absent on the empirical MSAs (left subfigure), as it reduces to zero. For simulated MSAs it is the true tree topology used for sequence simulation. The results in the right subfigure correspond to those already presented in the main text (Figure 5).

### Holdout-Validation: Supplementary results

Figure S12 presents the plausibility assessment results from the HV benchmarking, on empirical (left), and simulated (right) *long* MSAs. Results on empirical MSAs (left subfigure) have already been presented and discussed in the main text (Figure 4). Nonetheless, we also include them here for the sake of completeness. We do not report exact numerical results for simulated MSAs and confine ourselves to some general observations. The performance patterns are similar across empirical and simulated datasets, although the differences are less pronounced on simulated MSAs. Notably, the performance deterioration of HV versions compared to standard RAxML-NG (when restrained to the model-training MSA) is analogous to the decrease we observe on ES relative to standard RAxML-NG (both on the training MSA). This similarity possibly reflects the tree search heuristics of HV and ES, which are closely related (see Methods). Both heuristics appear to terminate prematurely when executed on a fraction of AA MSAs, as previously reported for ES (Togkousidis et al., 2025). Nonetheless, adopting this heuristic in HV constituted the only practical choice, since it follows the widely accepted holdout-based strategy used in deep learning (Montavon et al., 2012). Implementing a different holdout-based heuristic would have been difficult to theoretically justify. These issues warrant further investigation in the context of future work.
Figure S13 illustrates the average quartet distance distributions between the ML trees inferred by each version and the corresponding reference tree. For empirical MSAs, the reference tree is the ML tree inferred by standard RAxML-NG, on the training MSA of the corresponding MSA-split. Hence, the distribution of standard RAxML-NG (training MSAs) on empirical MSAs reduces to zero. For simulated MSAs, the reference tree is the true tree used for sequence simulation. The pairwise quartet distances, calculated between each ML tree inferred by each version, and the reference tree topology, are averaged over the 10 ML trees that each tool version infers per dataset. Results on simulated MSAs (right subfigure) have already been presented and discussed in the main text (Figure 5) but are also

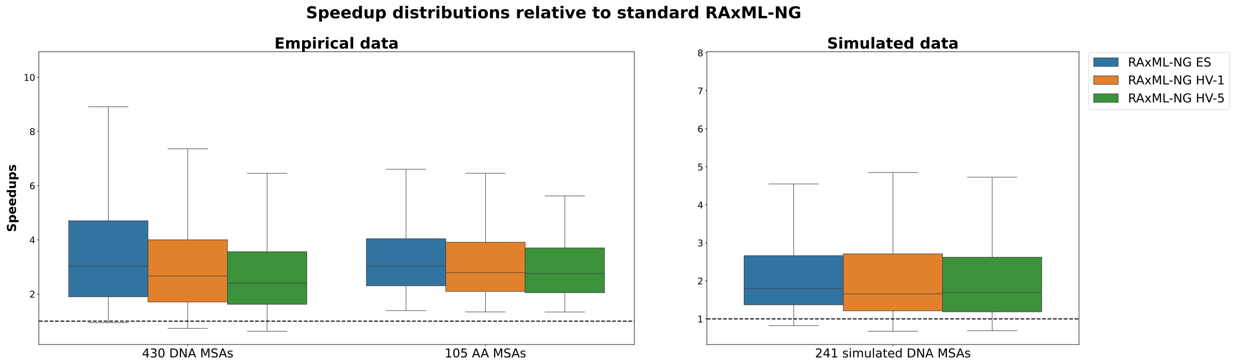


**Figure S14.** Speedup distributions of RAxML-NG ES and HV versions, relative to standard RAxML-NG inferences conducted on the training MSAs, for empirical (left subfigure) and simulated (right subfigure) MSAs. The dashed lines at the bottom of each subplot correspond to speedups of $1\times$.

included here for the sake of completeness.

On empirical DNA MSAs, the mean values of the average quartet distance distributions are: 0.014 for ES, 0.039 for both HV versions, and 0.033 for standard RAxML-NG using the model-training sites. The corresponding mean values for empirical AA MSAs are: 0.043 for ES, 0.058 for both HV versions, and 0.041 for standard RAxML-NG on model-training MSAs. Notably, although standard RAxML-NG (on model-training MSAs) outperforms ES on protein datasets in terms of mean values, the corresponding medians do favor ES. The median quartet distance for ES is 0.001, compared to 0.011 for RAxML-NG (model-training MSAs). This difference can also be observed by considering the shapes of the two distributions in Figure S13, since the ES distribution is skewed toward lower values. Thus, ES more frequently approximates the reference tree topology than RAxML-NG (model-training MSAs), albeit it also yields more extreme outliers. Most importantly, the results show that HV consistently underperforms relative to all other versions.

Finally, Figure S14 illustrates the speedup distributions of the ES and HV versions, relative to standard RAxML-NG execution times on the training MSA. We compute speedups by comparing the accumulated runtimes of each version for conducting 10 ML tree inferences. For empirical DNA MSAs, the average speedups are $3\times$ for HV-1, $2.7\times$ for HV-5, and $3.5\times$ for ES; on AA MSAs, the corresponding averages are $3.1\times$ for HV versions and $3.3\times$ for ES, respectively. On simulated DNA MSAs, HV versions show an average speedup of $2\times$, and ES a speedup of $2.2\times$. Overall, the HV versions do not yield any runtime improvement over ES on the tested datasets. Moreover, we should cautiously interpret the reported ES speedups, as the analyzed datasets show "easier" Pythia scores (see Figure S2). In our previous ES benchmark (Togkousidis et al., 2025), where we used a more representative distribution of MSAs with respect to Pythia scores, we obtained average speedups of $\sim 5\times$ for ES, relative to standard RAxML-NG, which is substantially higher compared to the speedups reported here. Thus, additional benchmarking on more challenging datasets is required to obtain conclusive results.
